# Exo-β-*N*-acetylmuramidase NamZ of *Bacillus subtilis* is the founding member of a family of exo-lytic peptidoglycan hexosaminidases

**DOI:** 10.1101/2021.01.10.425899

**Authors:** Maraike Müller, Matthew Calvert, Isabel Hottmann, Robert Maria Kluj, Tim Teufel, Katja Balbuchta, Alicia Engelbrecht, Khaled A. Selim, Qingping Xu, Marina Borisova, Alexander Titz, Christoph Mayer

## Abstract

Endo-β-*N*-acetylmuramidases, commonly known as lysozymes, are well-characterized antimicrobial enzymes that potentially lyse bacterial cells. They catalyze an endo-lytic cleavage of the peptidoglycan, the structural component of the bacterial cell wall; i.e. they hydrolyze glycosidic *N*-acetylmuramic acid (MurNAc)-β-1,4-*N*-acetylglucosamine (GlcNAc)-bonds within the heteroglycan backbone of peptidoglycan. In contrast, little is known about exo-β-*N*-acetylmuramidases, catalyzing an exo-lytic cleavage of β-1,4-MurNAc entities from the non-reducing ends of peptidoglycan chains. Such an enzyme was identified earlier in the bacterium *Bacillus subtilis*, but the corresponding gene has remained unknown so far. We identified *ybbC* of *B. subtilis,* renamed *namZ*, as encoding the reported exo-β-*N*-acetylmuramidase. A Δ*namZ* mutant accumulated specific cell wall fragments and showed growth defects under starvation conditions, indicating a role of NamZ in cell wall turnover. Recombinant NamZ protein specifically hydrolyzed the artificial substrate para-nitrophenyl β-MurNAc and the peptidoglycan-derived disaccharide MurNAc-β-1,4-GlcNAc. Together with the exo-β-*N*-acetylglucosaminidase NagZ and the exo-muramoyl-L-alanine amidase AmiE, NamZ degraded intact peptidoglycan by sequential hydrolysis from the non-reducing ends. NamZ is a member of the DUF1343 protein family of unknown function and shows no significant sequence identity with known glycosidases. A structural model of NamZ revealed a putative active site located in a cleft within the interface of two subdomains, one of which constituting a Rossmann-fold-like domain, unusual for glycosidases. On this basis, we propose that NamZ represents the founding member of a novel family of peptidoglycan hexosaminidases, which is mainly present in the phylum Bacteroidetes and, less frequently, within Firmicutes (Bacilli, Clostridia), Actinobacteria and Gammaproteobacteria.

## INTRODUCTION

The enzymatic degradation of carbohydrate polymers (glycans) in nature is generally mediated by endo- and exo-lytic glycosidases, which may act synergistically (1–3). Endo-glycosidases cleave glycosidic bonds within the inner part of glycan chains (endo-lytic cleavage), thereby solubilizing the polymers and providing oligomeric substrates for exo-glycosidases and exo-glucanases to act on, which subsequently chip off monomeric carbohydrates or disaccharides, respectively, from the non-reducing end or, in rare cases, from the reducing end (exo-lytic cleavage) (2, 4). Accordingly, cellulose (poly-β-1,4-glucose), the main glycan component of plants, and chitin (poly-β-1,4-*N*-acetylglucosamine), the main glycan component of fungi and arthropods, are degraded by various organisms involving complex sets of lytic enzymes. These include endo-lytic glycanases (endo-cellulases or endo-chitinases, respectively) and exo-glucanases, (cellobiohydrolases and diacetylchitobiohydrolases, respectively) as well as exo-lytic glycosidases (β-glucosidases/cellobiases and β-*N*-acetylglucosaminidases/diacetylchitobiases, respectively) (2, 4).

Considerably less is known about the synergistic cooperation of endo- and exo-lytic enzymes degrading peptidoglycan, the cell wall heteropolymer of bacteria. Peptidoglycan is composed of polysaccharide chains of alternating β-1,4-linked *N*-acetylglucosamine (GlcNAc) and *N*-acetylmuramic acid (MurNAc) carbohydrates that are interweaved forming a network via short oligopeptides linked to the 3-O-lactoyl residues of MurNAc (5). The lack of knowledge about the synergistic degradation of peptidoglycan is surprising, because endo-muramidases, catalyzing endo-lytic cleavage of the MurNAc-β-1,4-GlcNAc bond in the middle of the glycan backbone of the peptidoglycan, belong to the class of the best studied enzymes, notably hen’s egg white lysozyme, representing prominent antimicrobial defense molecules (6, 7). Most knowledge about peptidoglycan decomposition, however, relates to a bacterial metabolic process known as peptidoglycan turnover and recycling (8–10). During growth, bacteria need to degrade their own peptidoglycan to allow cell expansion, cell division and daughter cell separation. Besides peptidoglycan endopeptidases and amidases targeting the peptide part of the peptidoglycan, these turnover processes involve endo-lytic glycosidases that cleave either of the two glycosidic bonds within the peptidoglycan backbone: the MurNAc-β-1,4-GlcNAc-linkage (endo-β-*N*-acetylmuramidases; e.g. lysozymes) or the GlcNAc-β-1,4-MurNAc-linkage (endo-β-*N*-acetylglucosaminidases). The former enzymes include lysozyme-like muramidases, which catalyze a hydrolytic cleavage, and also a special type of muramidases, called lytic transglycosylases, which catalyze non-hydrolytic cleavage of the same bond, releasing products containing a 1,6-anhydro-glycosidic form of MurNAc (anhMurNAc) via an intramolecular transglycosylation reaction (11, 12). The latter enzymes are known as peptidoglycan-hydrolysing endo-β-*N*-acetylglucosaminidases (e.g. Atl^Glc^ or SagB of *S. aureus*; (13–15)). Whereas endo-β-N-acetylmuramidases and lytic transglycosylases generate oligosaccharides carrying a GlcNAc at the non-reducing end, endo-β-*N*-acetylglucosaminidases produce glycans with a MurNAc at the non-reducing end. Peptidoglycan fragments carrying a GlcNAc at the non-reducing end are further processed by exo-lytic β-*N*-acetylglucosaminidases (exo-GlcNAc’ases), which cleave off terminal, non-reducing GlcNAc residues. These enzymes are commonly found in various organisms and are classified in different families of glycosidases (e.g. CAZY families 3, 20, 84) [URL: https://www.cazypedia.org]. Many of them have been biochemically and mechanistically characterized in great detail, e.g. β-*N*-acetylglucosaminidases belonging to family 3 of glycosidases, such as NagZ of *Bacillus subtilis*, *Escherichia coli* and *Pseudomonas aeruginosa*, are well-known enzymes of the bacterial cell wall recycling pathways (16–18). In contrast, almost nothing is known about exo-lytic β-*N*-acetylmuramidases, which cleave terminal, non-reducing MurNAc residues that occur in peptidoglycan fragments generated by endo-β-*N*-acetylglucosaminidases. The only reports of an enzyme with exo-β-*N*-acetylmuramidase activity come from Del Rio and co-workers (19, 20). They identified such an enzyme entity occurs partly in the medium and partly bound to cells of *B. subtilis* at the end of exponential growth, and biochemically characterized the purified enzyme (19, 20). The encoding gene, however, has remained unknown, so far. We are investigating the function of peptidoglycan recycling and MurNAc catabolic operon of *B. subtilis* 168 (*murQ-murR-murP-amiE-nagZ-ybbC*) (16, 21). This operon consists of genes required for MurNAc catabolism: *murQ*, encoding a cytoplasmic MurNAc 6-phosphate (MurNAc-6P) lactyl ether hydrolase (22), *murR*, encoding a MurNAc-6P-sensing transcriptional regulator (23), and *murP*, encoding a MurNAc-specific phosphotransferase type transporter (24). Further, the operon contains *amiE* and *nagZ*, encoding two secreted exo-acting hydrolases, the exo-muramoyl-L-alanine amidase AmiE and the exo-β-*N*-acetylglucosaminidase NagZ, which are required for muropeptide rescue in *B. subtilis,* as shown earlier by our group (8, 16). We report here the identification of *ybbC,* the last gene within this operon, to encode the exo-*N*-acetylmuramidase of *B. subtilis* 168. The enzyme, referred to as NamZ hereafter, was biochemically characterized using the chromogenic substrate para-nitrophenyl 2-acetamido-3-O-(D-1-carboxyethyl)-2-deoxy-β-D-glucopyranoside (pNP-MurNAc) and the natural, peptidoglycan-derived disaccharide MurNAc-GlcNAc. NamZ specifically hydrolyzed pNP-MurNAc, MurNAc-GlcNAc and peptidoglycan fragments carrying a β-1,4-MurNAc entity at the non-reducing end, whereas the NagZ substrates pNP-GlcNAc, GlcNAc-MurNAc and peptidoglycan fragments carrying a β-1,4-GlcNAc entity at the non-reducing end are not hydrolyzed by NamZ. NamZ together with the exo-muramoyl-L-alanine amidase AmiE and the exo-*N*-acetylglucosaminidase NagZ of *B. subtilis* are able to cleave peptidoglycan and fragments thereof (muropeptides), sequentially from the non-reducing end, thereby releasing MurNAc, GlcNAc and peptidoglycan peptides, which can be further metabolized (16, 21). Thus, in contrast to the degradative systems of the homopolymers cellulose and chitin, the cleavage of the heteropolymer peptidoglycan involves two distinct sets of exo- and endo-lytic β-hexosaminidases: exo- and endo-acting β-*N*-acetylglucosaminidases as well as exo- and endo-acting β-*N*-acetylmuramidases. Here we propose that NamZ of *B. subtilis* is the founding member of a unique family of exo-lytic β-*N*-acetylmuramidases, which is mainly present in the phylum of *Bacteriodetes* and, less frequently, within *Bacilli*, *Clostridia*, *Actinobacteria* and *Gammaproteobacteria*.

## RESULTS

### The B. subtilis ybbC (namZ) mutant displays defects in cell growth and exo-β-N-acetylmuramidase function

The function of *ybbC,* the last gene within the peptidoglycan/MurNAc recycling operon of *B. subtilis* 168 (*murQ-murR-murP-amiE-nagZ-ybbC*) remained unknown so far (16, 21). It encodes a putative enzyme with no significant sequence identity towards any protein of known function, containing a conserved protein domain of unknown function (PFAM DUF1343 or PF07075). The location of *ybbC* in an operon next to *nagZ* and transcriptional analyses of *B. subtilis* cells grown under different conditions revealed that the expression of both genes is co-regulated (16, 25). We hypothesized that *ybbC* may encode the enigmatic, secreted exo-β-*N*-acetylmuramidase, identified by Del Rio *et al.*, and functioning in conjunction with NagZ (16,19,20). A putative signal peptide motif (MRKTIFAFLTGLMMFGTITAASA/ SPD; /, cleavage site according to signal P-5.0 (26)) suggested that the *ybbC*-encoded protein is secreted. To obtain a first indication that *ybbC* encodes the unknown exo-β-*<u>N</u>*-<u>a</u>cetyl<u>m</u>uramidase coregulated with Nag<u>Z</u>, termed NamZ, we compared growth and extracellular accumulation of cell wall fragments in rich medium (LB) of *B. subtilis* 168 (wild-type; WT), Δ*nagZ* and Δ*namZ* (Δ*nagZ::erm* and Δ*namZ::erm* obtained from the Bacillus Genetic Stock Center (27) were rendered markerless by excision of the erythromycin resistance cassettes, as described in the Experimental Procedures). We observed that the two mutants showed a similar growth behavior, which differed from that of the WT, in that the two mutants reached lower OD_600_ values in early stationary phase compared to the WT strain (Supporting Figure S1). This is consistent with the reported expression profile of *nagZ,* which protein levels were very low during the exponential phase, but strongly increased during transition and stationary phase (16). However, following prolonged stationary phase (52 h of growth; Supporting Figure S1), both the Δ*namZ* and the Δ*nagZ* mutant displayed a diminished lytic phenotype compared to WT, as reported earlier for *nagZ* (16).

A mass spectrometric analysis of the supernatants of mutants and WT cells was conducted to identify the cell wall fragments that specifically accumulated in the supernatants of Δ*namZ* and Δ*nagZ*, and may indicate the absence of exo-muramidase function in the *namZ* deficient strain. As expected, no significant accumulation of small soluble products was observed in the supernatants of exponentially growing cells (5h; data not shown). Further, the analysis of stationary phase supernatants (20h) also revealed only minor differences in the accumulation of small extracellular cell wall fragments. A compound with a mass-to-charge ratio in positive ion mode of (M+H)^+^ 497.204 m/z, corresponding to MurNAc-GlcNAc (theoretical (M+H)^+^ 497.1977 m/z; Supporting Table S1), which was expected to accumulate in the supernatants of Δ*namZ* mutants, was found in only very tiny amounts, albeit about two-fold more in the Δ*namZ* that in the WT stain (Figure 1). Rather, another compound, which was identified as the trisaccharide MurNAc-GlcNAc-1,6-anhydroMurNAc (MurNAc-GlcNAc-anhMurNAc) by its mass-to-charge ratio (M+H)^+^ of 754.284 m/z (theoretical (M+H)^+^ 754.2877 m/z; Supporting Table S1), accumulated in considerable amounts and about two-fold more compared to WT (Figure 1). This product of peptidoglycan cleavage by a lytic transglycosylase, is expected to accumulate in a strain defective for an exo-lytic *N*-acetylmuramidase. This trisaccharide was absent in the Δ*nagZ* mutant, which instead accumulated large amounts of GlcNAc-anhMurNAc and to a lesser extent also GlcNAc-MurNAc. Both compounds were unequivocally identified by their (M+H)^+^ of 479.185 m/z and 497.196 m/z, respectively (Figure 1), and by *in vitro* cleavage with the recombinant NagZ enzyme (data not shown). GlcNAc-anhMurNAc and GlcNAc-MurNAc disaccharides did not accumulate in the Δ*namZ* mutant and the WT strain, which indicates that NagZ is responsible for cleaving GlcNAc-anhMurNAc/GlcNAc-MurNAc. Consistently, an accumulation of anhMurNAc ((M+H)^+^ of 276.106 m/z) was observed in the supernatant of WT and Δ*namZ* strains but was absent in Δ*nagZ.* Although the amount of potential NamZ substrates that accumulated in the supernatant of a *B. subtilis* Δ*namZ* mutant was surprisingly low, the identity of the small soluble peptidoglycan fragments with a MurNAc entity at the reducing end supports our hypothesis that *namZ* encodes a secreted exo-*N*-acetylmuramidase.

**Figure 1.**
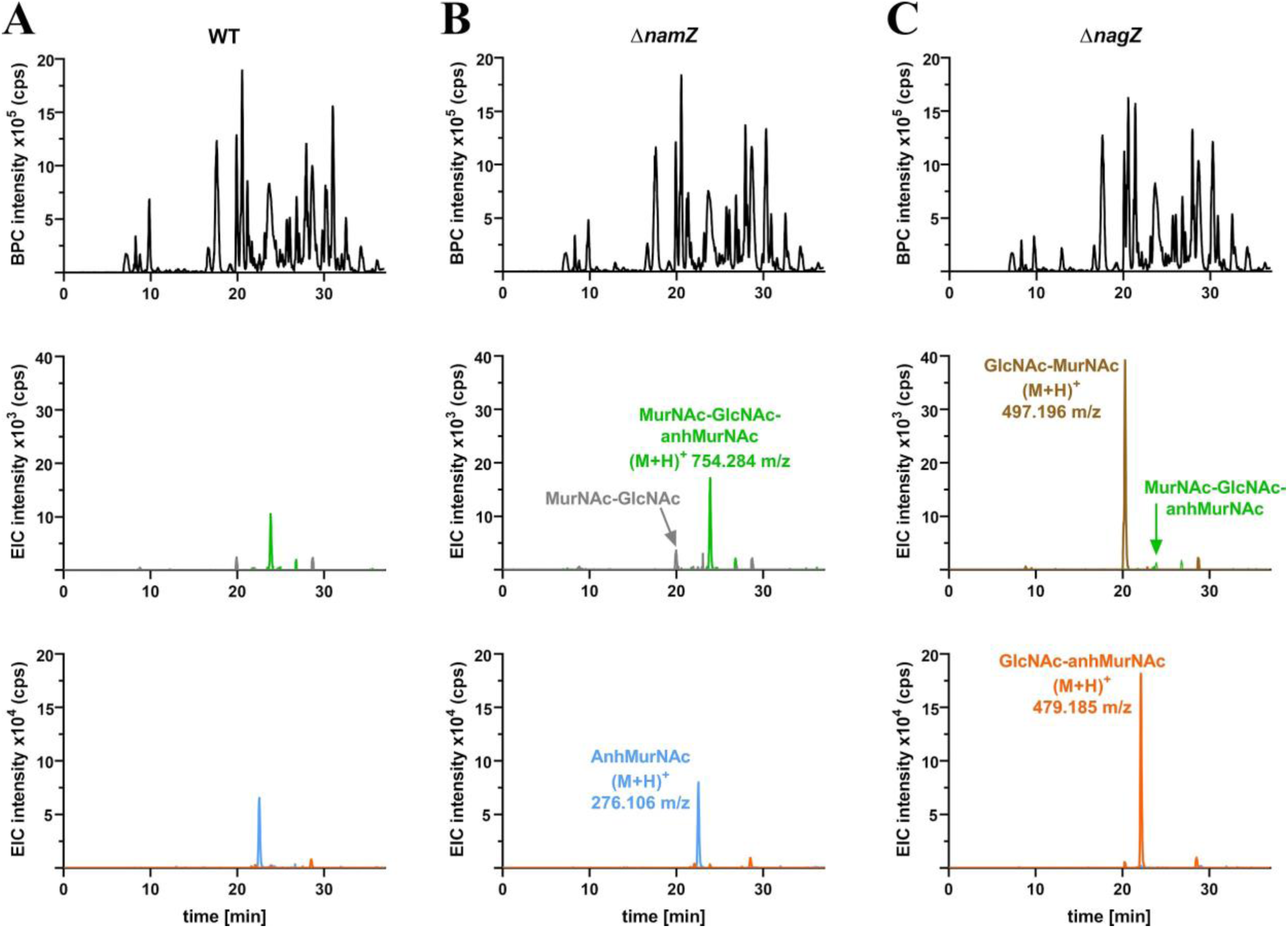
Accumulation of specific cell wall fragments in growth supernatants of *B. subtilis* WT, Δ*namZ* and Δ*nagZ* cells. Supernatants of stationary phase cultures (grown for 20h in LB) of *B. subtilis* WT **(A)**, Δ*namZ* **(B)** and Δ*nagZ* **(C)**, were analyzed by LC-MS for the accumulation of small cell wall fragments. Shown are the base peak chromatograms (BPCs; black) and extracted ion chromatograms (EIC) for the disaccharides MurNAc-GlcNAc (grey), GlcNAc-MurNAc (brown), GlcNAc-anhMurNAc (orange) as well as anhMurNAc (light blue) and the trisaccharide MurNAc-GlcNAc-anhMurNAc (green). To unequivocally identify the accumulation products, these were treated with recombinant NagZ (8).

**Table 1.**
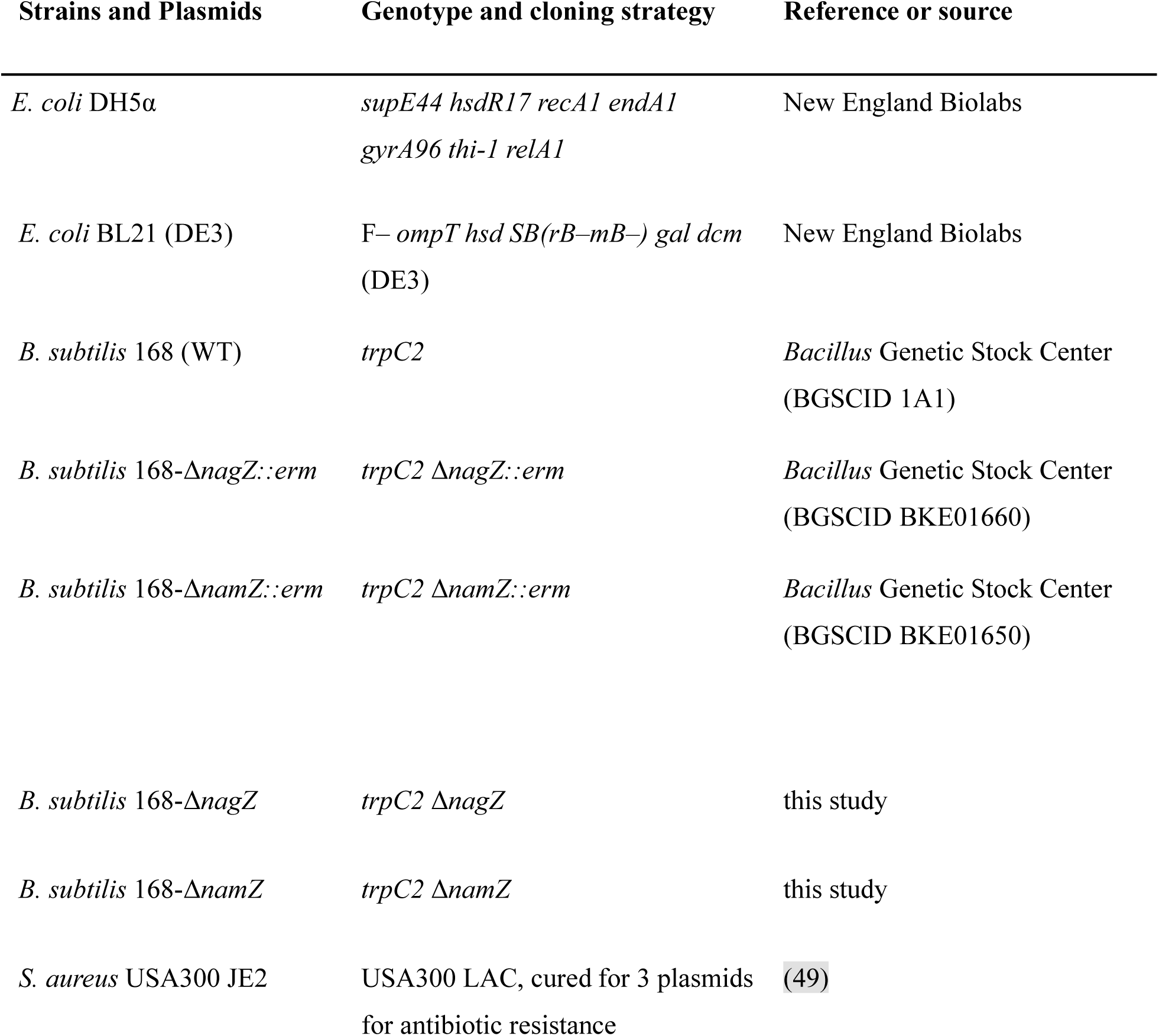
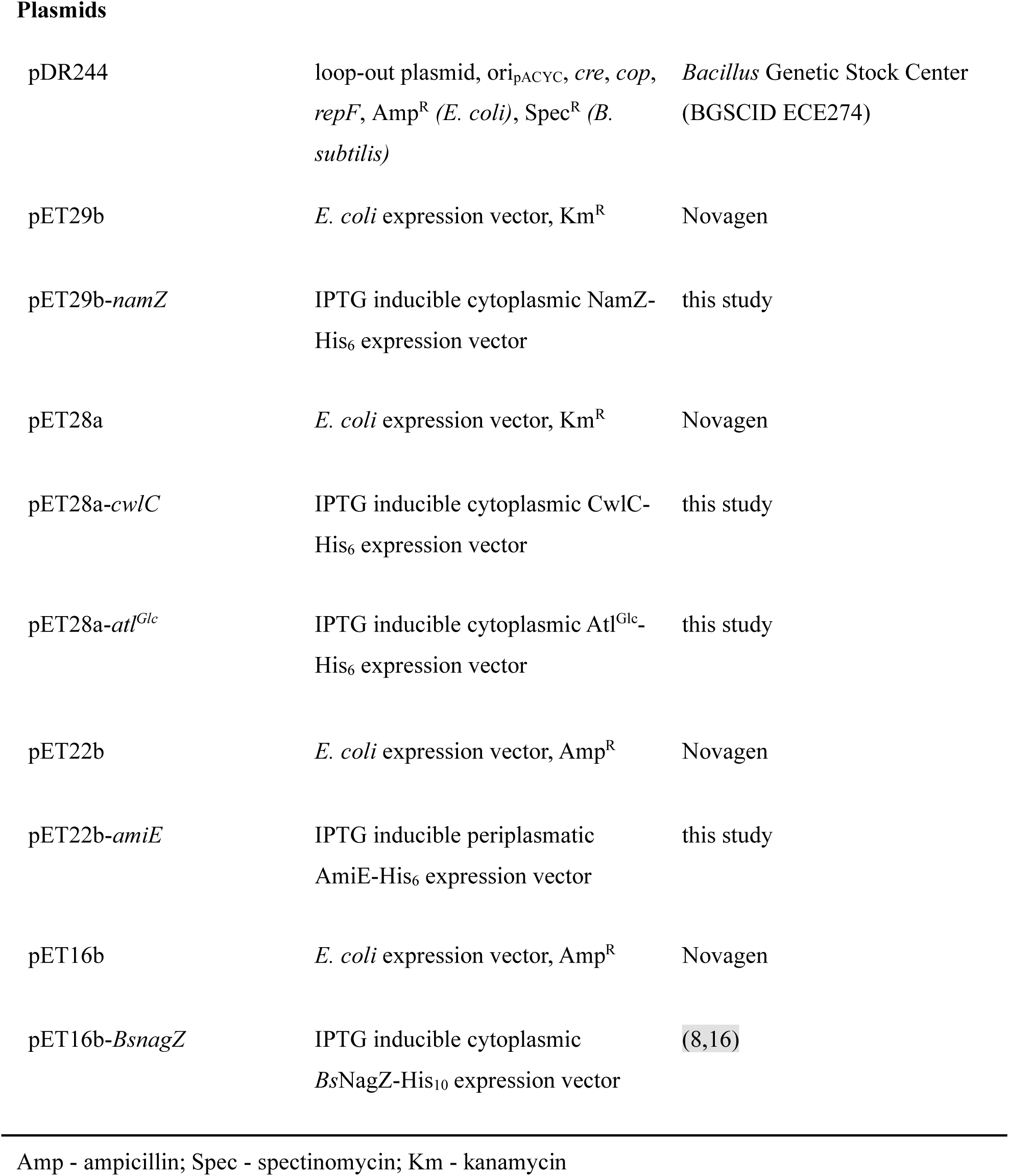
Strains and plasmids.

| Strains and Plasmids | Genotype and cloning strategy | Reference or source |
| --- | --- | --- |
| *E. coli* DH5α | *supE44 hsdR17 recA1 endA1*<br>*gyrA96 thi-1 relA1* | New England Biolabs |
| *E. coli* BL21 (DE3) | F– *ompT hsd SB(rB–mB–) gal dcm*<br>(DE3) | New England Biolabs |
| *B. subtilis* 168 (WT) | *trpC2* | *Bacillus* Genetic Stock Center<br>(BGSCID 1A1) |
| *B. subtilis* 168-Δ*nagZ::erm* | *trpC2* Δ*nagZ::erm* | *Bacillus* Genetic Stock Center<br>(BGSCID BKE01660) |
| *B. subtilis* 168-Δ*namZ::erm* | *trpC2* Δ*namZ::erm* | *Bacillus* Genetic Stock Center<br>(BGSCID BKE01650) |
| *B. subtilis* 168-Δ*nagZ* | *trpC2* Δ*nagZ* | this study |
| *B. subtilis* 168-Δ*namZ* | *trpC2* Δ*namZ* | this study |
| *S. aureus* USA300 JE2 | USA300 LAC, cured for 3 plasmids<br>for antibiotic resistance | (49) |
| Plasmids |  |  |
| pDR244 | loop-out plasmid, ori<sub>pACYC</sub>, *cre*, *cop*, *repF*, Amp<sup>R</sup> (*E. coli*), Spec<sup>R</sup> (*B. subtilis*) | *Bacillus* Genetic Stock Center (BGSCID ECE274) |
| pET29b | *E. coli* expression vector, Km<sup>R</sup> | Novagen |
| pET29b-*namZ* | IPTG inducible cytoplasmic NamZ-His<sub>6</sub> expression vector | this study |
| pET28a | *E. coli* expression vector, Km<sup>R</sup> | Novagen |
| pET28a-*cwlC* | IPTG inducible cytoplasmic CwlC-His<sub>6</sub> expression vector | this study |
| pET28a-*atl*<sup>Glc</sup> | IPTG inducible cytoplasmic Atl<sup>Glc</sup>-His<sub>6</sub> expression vector | this study |
| pET22b | *E. coli* expression vector, Amp<sup>R</sup> | Novagen |
| pET22b-*amiE* | IPTG inducible periplasmatic AmiE-His<sub>6</sub> expression vector | this study |
| pET16b | *E. coli* expression vector, Amp<sup>R</sup> | Novagen |
| pET16b-*BsnagZ* | IPTG inducible cytoplasmic BsnagZ-His<sub>10</sub> expression vector | (8,16) |

Amp - ampicillin; Spec - spectinomycin; Km - kanamycin

### Recombinant NamZ forms a dimer in solution

To further characterize NamZ, the enzyme was cloned and heterologously expressed in *E. coli* as C-terminal His_6_-tag fusion protein, and subsequently purified to apparent homogeneity via Ni^2+^ affinity chromatography followed by size exclusion chromatography, as revealed by sodium dodecyl sulfate-polyacrylamide gel electrophoresis (SDS-PAGE) analysis (Supporting Figure S2). SDS-PAGE analysis of NamZ under denaturing conditions revealed the monomeric protein, in agreement with the calculated mass of 44.77 kDa. Del Rio and colleagues reported that the exo-*N*-acetylmuramidase entity is an enzyme with an apparent molecular mass of 90 kDa (20), suggesting that the enzyme forms a dimer in solution (Supporting Figure S2). Therefore, we determined the oligomeric state of NamZ in solution. The purified recombinant NamZ protein was subject to size exclusion chromatography coupled with multiangle light scattering (SEC-MALS). At the peak maximum, NamZ eluted clearly with an apparent mass according to MALS analysis of 89.3 ± 7.4 kDa in three independent experiments (Supporting Figure S2), confirming the expected theoretical mass of 89.54 kDa of the NamZ dimer. Based on SEC-MALS analysis, we concluded that the NamZ protein indeed appears mainly dimeric in solution, consistent with the previous observation (20).

### Identification of NamZ as a specific exo-N-acetylmuramidase using synthetic substrate pNP-MurNAc

Chromogenic and fluorogenic substrates greatly facilitate the characterization of glycosidases, as they allow the determination of enzymatic activity and specificity with high sensitivity; e.g. the β-galactosidase LacZ of *E. coli* is routinely assayed using ortho-nitrophenyl β-galactoypranoside (oNP-Gal) in the so-called β-Gal assay (28). Similarly, the activity of NagZ of *B. subtilis* had been assayed using chromogenic para-nitrophenyl β-*N*-acetylglucosaminide (pNP-GlcNAc) or, alternatively, the fluorogenic 4-methylumbelliferyl β-*N*-acetylglucosaminide (4MU-GlcNAc) (8,29,30). Del Rio *et al*. reported the synthesis and use of the fluorogenic substrate 4MU-MurNAc for the detection of exo-β-*N*-acetylmuramidase activity in the supernatant of *B. subtilis* cultures (20, 31). We decided to synthesize the analogous chromogenic substrate para-nitrophenyl β-*N*-acetylmuramic acid (pNP-MurNAc; para-nitrophenyl 2-acetamido-3-O-(D-l-carboxyethyl)-2-deoxy-β-D-glucopyranoside), starting from commercially available pNP-GlcNAc in analogy to a previously published procedure (32). In brief, pNP-GlcNAc was transformed into its 4,6-benzylidene derivative followed by alkylation of the 3-OH group using rac-2-chloropropionic acid. Direct methylation of the carboxyl group allowed purification and separation of the desired diastereomer from the minor *iso*-MurNAc diastereomer. Final deprotection under acidic conditions for the removal of the benzylidene group and basic conditions for methyl ester cleavage resulted in 67% yield of pNP-MurNAc. A similar procedure was followed to isolate pNP-iso-MurNAc derivative (see Supporting Information for details). Exo-β-*N*-acetylmuramidase activity of recombinant NamZ was demonstrated by the release of nitrophenolate from the chromogenic substrate pNP-MurNAc (Figure 2). To determine the specificity of NamZ, the chromogenic substrates pNP-GlcNAc and oNP-GalNAc were also tested. NamZ specifically hydrolyzed pNP-MurNAc, but not the other tested chromogenic substrates, which in turn were specifically cleaved by LacZ and NagZ. This assay identified NamZ as a highly specific exo-β-*N*-acetylmuramidase (Figure 2).

**Figure 2.**
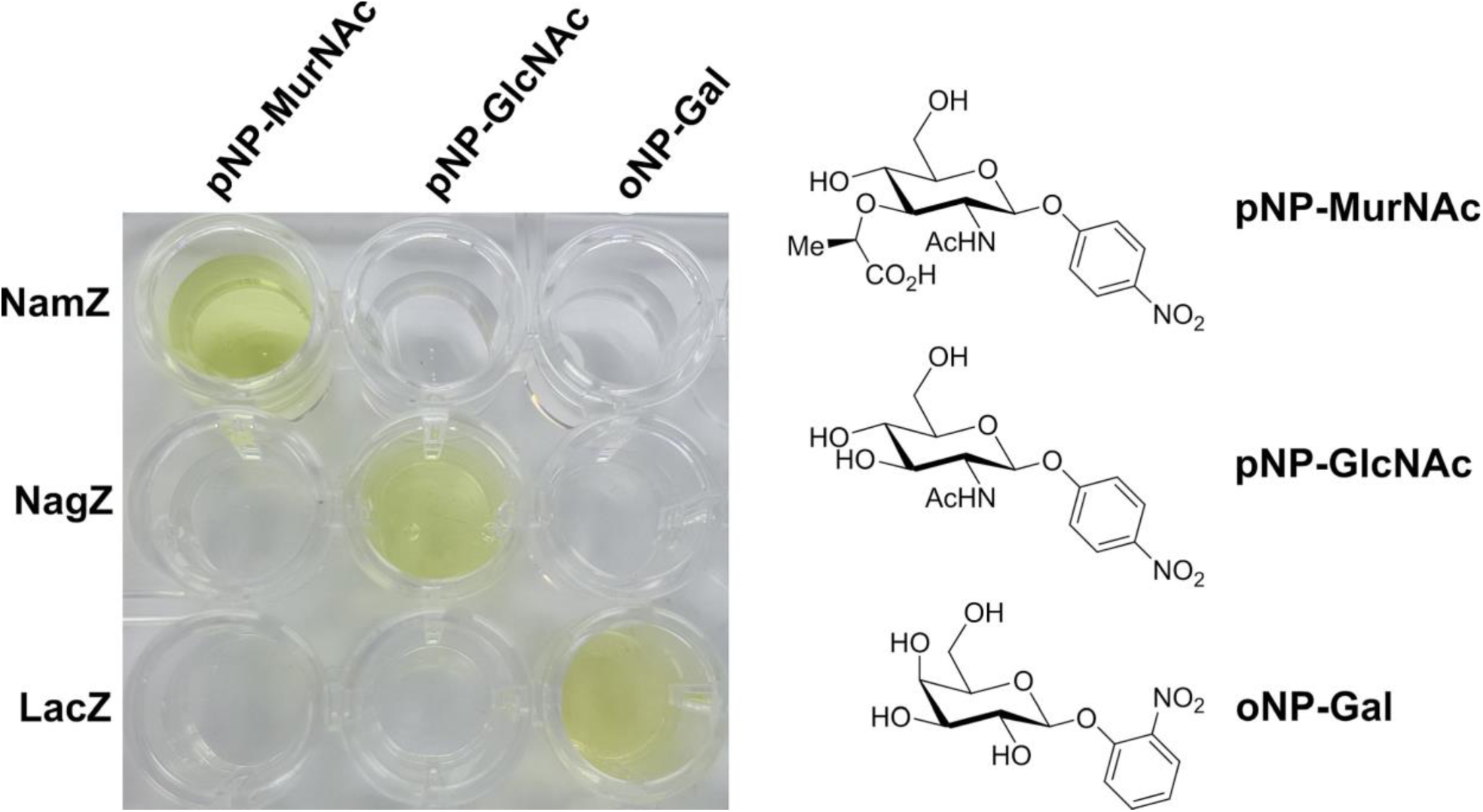
NamZ of *B. subtilis* specifically cleaves the chromogenic substrate pNP-MurNAc. The other tested chromogenic substrates, pNP-GlcNAc and oNP-Gal were not cleaved by NamZ, but are substrates of the exo-β-*N*-acetylglucosaminidase of *B. subtilis* (NagZ) and the β-D-galactosidase of *E. coli* (LacZ), respectively. Purified recombinant NamZ and NagZ from *B. subtilis*, as well as commercial LacZ (100 μg/ml each) were incubated for 30 min with the indicated chromogenic compound (100 μM each). Color intensity was enhanced by adding an equal volume of sodium borate buffer (100 mM, pH 10).

### Biochemical characterization of NamZ using pNP-MurNAc

Prior to the biochemical characterization of NamZ using pNP-MurNAc, we checked the purity of the synthetic substrate with HPLC-MS and NMR. The ^1^H- and ^13^C-NMR spectra of the synthetic pNP-MurNAc and pNP-isoMurNAc products indicate that both compounds are of high purity (> 95%) (for a detailed description of the synthesis see Supporting Information). In the methanolic stock solution of pNP-MurNAc used for enzyme reactions, the main compounds in the base peak chromatogram (BPC) observed by MS analysis showed a mass of (M+H)^+^ 415.132 m/z (retention time of 36.7 min) corresponding to the theoretical mass of pNP-MurNAc ((M+H)^+^ 415.1347 m/z; cf. Supporting Table S1). Occasionally we found an additional small peak at a retention time of 40.3 min with a mass of (M+H)^+^ 429.147 m/z, identified as methylated pNP-MurNAc (theoretical mass (M+H)^+^ 429.1504 m/z; cf. Supporting Table S1), which may have resulted from esterification of the acidic pNP-MurNAc in the prepared stock solution in MeOH. Since traces of the methylated pNP-MurNAc may inhibit the NamZ reaction, the methylation was removed by pre-incubation of the pNP-MurNAc substrate for 30 minutes at pH 8.0 prior to the NamZ reactions.

We first determined stability and reaction optima of recombinant NamZ at temperatures between 4 and 60 °C and pH values between 2 and 10, using the chromogenic substrate pNP-MurNAc. Stability was measured by the loss of activity of NamZ after 30 min of pre-incubation at different temperatures or pH conditions and the optima were determined by direct reactions of NamZ at different temperature or pH conditions. NamZ was shown to be stable from 4 °C to 37 °C, with a short-term maximal activity at around 37 °C (Supporting Figure S3). NamZ was also shown to be mostly stable over a pH range from 6.0 to 10.0, with a short-term maximal activity between pH 6.0 and 8.0 (Supporting Figure S3). Furthermore, NamZ retained its activity for several months after purification when stored at either −20 °C or 4 °C. We chose to determine the kinetic parameters of NamZ at 37 °C and in the pH-range between 7.0 to 8.0. Kinetic parameters for the chromogenic substrate were calculated using a standard curve for para-nitrophenol (Figure 3). The maximal velocity of the reaction of NamZ with the chromogenic substrate pNP-MurNAc was determined as *v*_max_ of 1.45 µmol min^-1^ mg^-1^ and a *K*_M_ of 0.125 mM at 37°C and pH 8.0, since the slightly basic pH increases the sensitivity in an continuous colorimetric assay.

**Figure 3.**
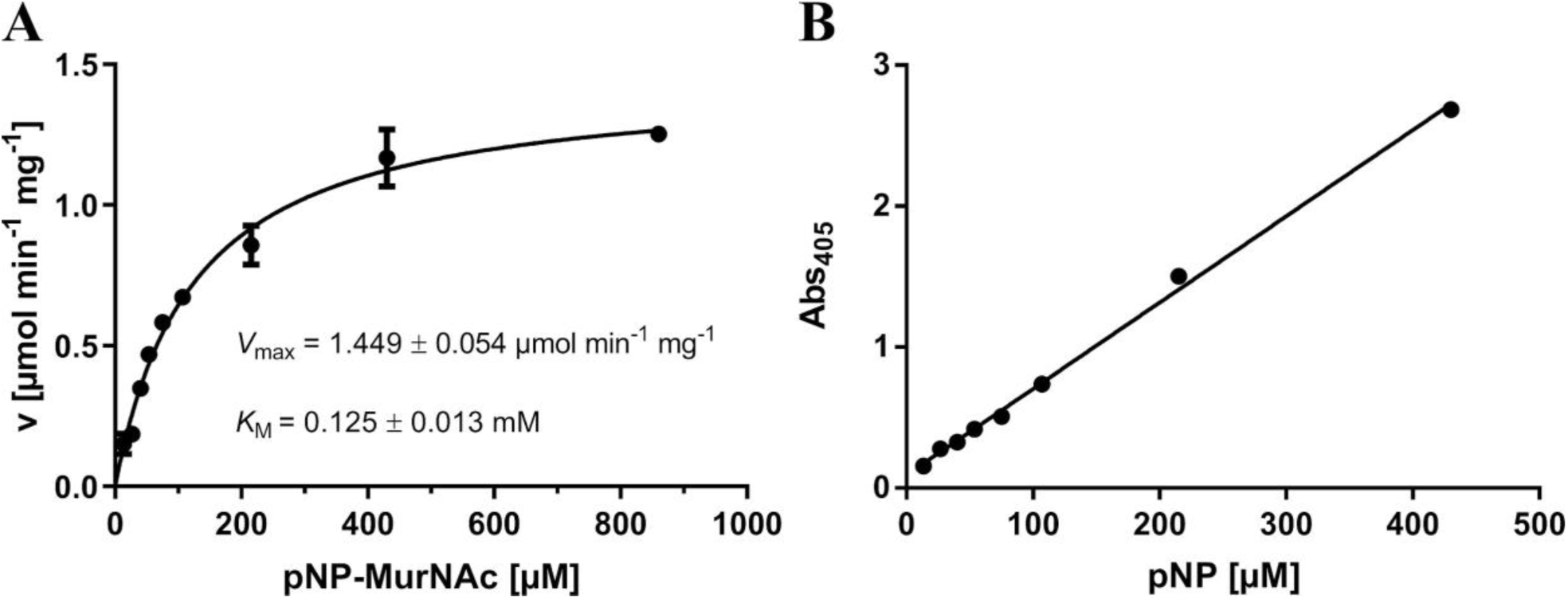
Determination of the kinetic parameters of NamZ using the chromogenic substrate pNP-MurNAc at pH 8.0. **(A)** Kinetic parameters using the chromogenic substrate pNP-MurNAc were determined as described in Experimental Procedures. **(B)** Standard curve for 4-nitrophenol (13 to 430 µM) was determined in 0.2 M phosphate buffer (pH 8.0). Absorption of pNP was measured at 405 nm. For all experiments, standard errors (SEM) are indicated and calculated out of three biological replicates.

### Preparation of MurNAc-GlcNAc from peptidoglycan enzymatic digests

We aimed to biochemically characterize NamZ further, using the disaccharide MurNAc-GlcNAc, the proposed minimal natural substrate. To prepare MurNAc-GlcNAc, peptidoglycan isolated from *B. subtilis* was digested with recombinant muramyl-L-alanine amidase (CwlC of *B. subtilis*) and endo-*N*-acetylglucosaminidase (Atl^Glc^ of *S. aureus*) as described elsewhere (15, 33). The crude enzyme digest contained large amounts of the disaccharide MurNAc-GlcNAc, as revealed by LC-MS analysis, but also crosslinked, non-crosslinked peptides and other saccharides in minor amounts. Therefore, the disaccharide was purified using semi-preparative HPLC, yielding a MurNAc-GlcNAc preparation that was mostly free of tri- and tetrapeptides as well as crosslinked tri-tetrapeptides as shown by LC-MS analysis (Supplementary Figure S4). The mass spectrum of the MurNAc-GlcNAc pool, appearing at retention time of 21.2 min, contained solely the molecule mass (M+H)^+^ 497.196 m/z, confirming that the MurNAc-GlcNAc preparation was of sufficient purity (Supporting Figure S4C).

### Biochemical characterization of NamZ using MurNAc-GlcNAc

Using the purified natural minimal substrate MurNAc-GlcNAc, NamZ activity was characterized. The disaccharide ((M+H)^+^ 497.196 m/z, retention time of 21.1 min) completely disappeared by adding recombinant NamZ (Figure S5A) and two new peaks appeared, which were identified as MurNAc (measured mass (M+H)^+^ 294.118 m/z, retention time of 21.8 min) and GlcNAc (measured mass (M+H)^+^ 222.097 m/z, retention time of 10.0 min) (Figure S5B).

NamZ requires a MurNAc entity at the non-reducing end, since GlcNAc-MurNAc was not cleaved as shown by LC-MS (determined (M+H)^+^ 497.198 m/z, retention time of 21.8 min) (Supporting Figure S6). In contrast, NagZ readily cleaved GlcNAc-MurNAc but not MurNAc-GlcNAc. After incubation with NagZ, the GlcNAc-MurNAc peak completely disappeared and two new peaks appeared that correspond to GlcNAc (measured mass (M+H)^+^ 222.097 m/z, retention time of 10.0 min) and MurNAc (measured mass (M+H)^+^ 294.118 m/z, retention time of 21.8 min) (Supporting Figure S6).

To determine the kinetic parameters of NamZ with MurNAc-GlcNAc as a substrate we chose a pH of 7.0 for the assay. For MurNAc-GlcNAc, the release of MurNAc by the action of NamZ was analyzed using HPLC-MS. Therefore, a standard curve for MurNAc (Figure 4) was created and kinetic parameters for the natural minimal substrate were calculated (Figure 4). The reaction of NamZ with MurNAc-GlcNAc as a substrate at pH 7.0 was much faster than with the chromogenic substrate pNP-MurNAc at pH 8.0 (*v*_max_ of 88.64 and 1.45 µmol min^-1^ mg^-1^), but the *K*_M_ for MurNAc-GlcNAc was a 30-fold higher than that for pNP-MurNAc (3.59 and 0.125 mM). With MurNAc-GlcNAc as a substrate, a first saturation was reached at a substrate concentration of 0.2 mM substrate. After this saturation, the curve increased again until the second saturation at a substrate concentration of 6 mM was reached. Because of these two saturations, the kinetic parameters for 0.2 mM MurNAc-GlcNAc as highest concentration were also calculated. Hence, a *v*_max_ of 5.99 µmol min^-1^ mg^-1^ and a *K*_M_ of 0.22 mM were determined (Supporting Figure S7).

**Figure 4.**
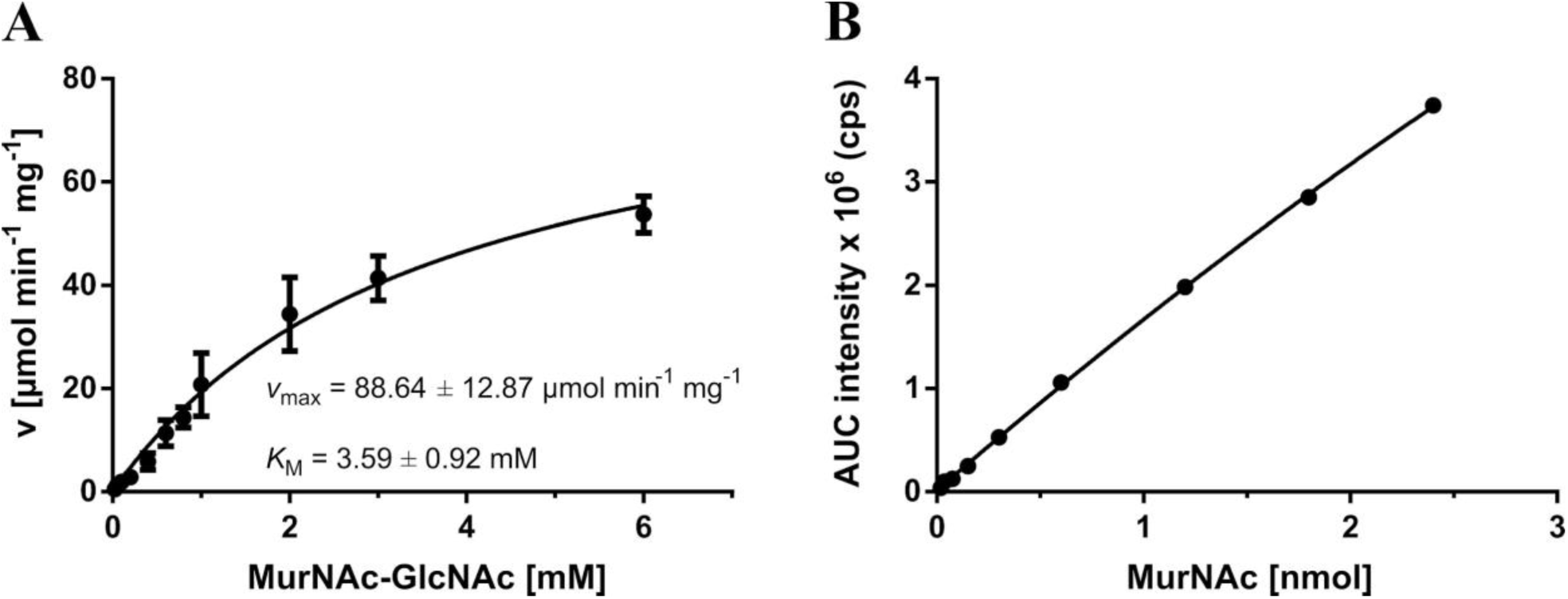
Determination of the kinetic parameters of NamZ using MurNAc-GlcNAc substrate at pH 7.0. **(A)** Kinetic parameters using purified MurNAc-GlcNAc were determined. Standard errors (SEM) are indicated and calculated out of three biological replicates. **(B)** Standard curve for MurNAc (0.019 to 2.4 nmol/3µl) was determined in 0.2 M phosphate buffer (pH 7.0) and stopping buffer (1% formic acid, 0.5% ammonium formate, pH 3.2) in equal amounts in a total volume of 50 µl. Samples were analyzed using HPLC-MS and the areas under the curve (AUCs) were generated using the extracted ion chromatograms (EICs) intensities for MurNAc ((M-H)^-^ 292.113 m/z).

### Sequential digest of peptidoglycan by NagZ, AmiE, and NamZ

*B. subtilis* 168 peptidoglycan was digested with NagZ, AmiE and NamZ in one reaction. To this end, peptidoglycan, the exo-β-*N*-acetylglucosaminidase NagZ, the *N*-acetylmuramyl-L-alanine amidase AmiE and NamZ were purified as described in the Experimental Procedures. Peptidoglycan was incubated with all three enzymes overnight and the generated products were analyzed using HPLC-MS in negative and positive ion mode. Analysis of degraded peptidoglycan revealed single cell wall carbohydrates and peptides (Figure 5).

**Figure 5.**
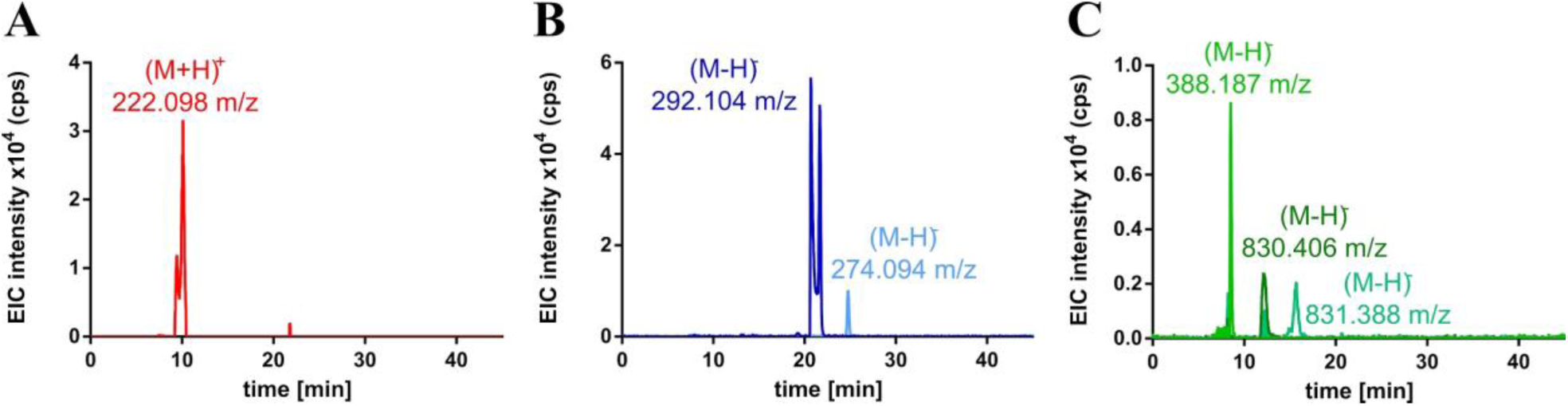
Sequential digest of peptidoglycan by NagZ, NamZ and AmiE. Purified peptidoglycan was sequentially digested by the action of NagZ, NamZ and AmiE and analyzed by HPLC-MS. Main products are shown as extracted ion chromatograms (EIC). Single cell wall carbohydrates could be identified with observed masses **(A)** (M+H)^+^ 222.098 m/z for GlcNAc (red) with a retention time of 10.1 min, **(B)** (M-H)^-^ 292.104 m/z for MurNAc (dark blue) with a retention time of 21.7 min and (M-H)^-^ 274.094 m/z for anhMurNAc (light blue) with a retention time of 24.8 min. **(C)** Peptides could be identified with observed masses (M-H)^-^ 388.187 m/z for tripeptide with an amidation (light green) with a retention time of 8.5 min, (M-H)^-^ 830.406 m/z for tri-tetrapeptide with two amidations (dark green) with a retention time of 12.1 min and (M-H)^-^ 831.388 m/z for tri-tetrapeptide with one amidation (turquoise) with a retention time of 15.1 min.

We could identify GlcNAc with an observed mass of (M+H)^+^ 222.098 m/z and a retention time of 10.1 min, MurNAc with an observed mass of (M-H)^-^ 292.104 m/z and a retention time of 21.7 min and anhMurNAc with an observed mass of (M-H)^-^ 274.094 m/z and a retention time of 24.8 min as single cell wall carbohydrates. Furthermore, peptides were identified as tripeptide with an amidation with an observed mass of (M-H)^-^ 388.187 m/z and a retention time of 8.5 min, cross-linked tri-tetrapeptide with two amidations with an observed mass of (M-H)^-^ 830.406 m/z and a retention time of 12.1 min and cross-linked tri-tetrapeptide with one amidation with an observed mass of (M-H)^-^ 831.388 m/z and a retention time of 15.1 min. Additionally, the mass spectra of the carbohydrates and peptide peaks were analyzed (Supporting Figure S8). The GlcNAc peak at the retention time of 10.1 min was identified by its m/z corresponding to GlcNAc (M+H)^+^ 222.098 m/z and the detection of m/z of the sodium adduct (M+H)^+^ 244.079 m/z and the water elimination product (M+H)^+^ 204.087 m/z as well as a product in which two water eliminations occured (M+H)^+^ 186.077 m/z. Further analysis of the MurNAc peak at a retention time of 21.7 min was identified by the m/z corresponding to MurNAc (M-H)^-^ 292.104 m/z as well as the MurNAc dimer ((M-H)^-^ 585.214 m/z). The anhMurNAc at a retention time of 24.8 min was identified by the m/z corresponding to anhMurNAc (M-H)^-^ 274.094 m/z, as well as the anhMurNAc dimer ((M-H)^-^ 549.197 m/z). Peptides released from the peptidoglycan were identified as the tripeptide (Ala-Glu-DAP) with one amidation with a retention time of 8.5 min, tri-tetrapeptide with two amidations at a retention time of 12.1 min and tri-tetrapeptide with one amidation at a retention time of 15.1 min. When peptidoglycan was digested with one of the three enzymes alone, only low amounts of cell wall monosaccharides and peptides could be identified. Hence, the joint and coordinated activity of all three enzymes NagZ, AmiE and NamZ, is needed to digest the peptidoglycan, thereby releasing the respective peptidoglycan-derived monosaccharides and peptides.

### Structure model of NamZ reveals unique glycosidase fold

NamZ shares significant homology with two proteins from *Bacteroides fragilis* NCTC 9343, BF0379 (40% sequence identity, PDB ID 4jja) and BF0371 (44% sequential identity, PDB ID 4k05), the structures of which were determined and deposited in the Protein Data Bank (Joint Center for Structural Genomics). Hitherto, nothing is known about the biological function of these proteins. We built a homology model of NamZ based on these structures, using HHPred (34) (Figure 6). Even though NamZ shares higher overall sequence homology with BF0371, the residues around the putative active site of NamZ and BF0379 are more conserved, suggesting that BF0379 and NamZ may share a very similar function. NamZ consists of two domains, an N-terminal catalytic domain (34–258) and a C-terminal auxiliary domain (259–414). The N-terminal catalytic domain has an α/β/α three-layer sandwich architecture with a central parallel β-sheet topology of 321456 (order of β-strands in the sheet). A putative catalytic residue (Glu92) is located at the C-terminal end of the second β-strand. The C-terminal auxiliary domain has an α/β architecture with 3 strands. A putative active site cavity is formed between the domain interface defined by both domains. A positive charged arginine (Arg279) from this C-terminal domain points toward the cavity. Surprisingly, the catalytic domain has a Rossmann-like fold, which is usually involved in binding nucleotides. However, the catalytic domain of NamZ does not process the glycine-rich signature nucleotide binding motif. Moreover, the nucleotide binding region is partially occupied by its C-terminal region of the catalytic domain (residues 245-251).

**Figure 6.**
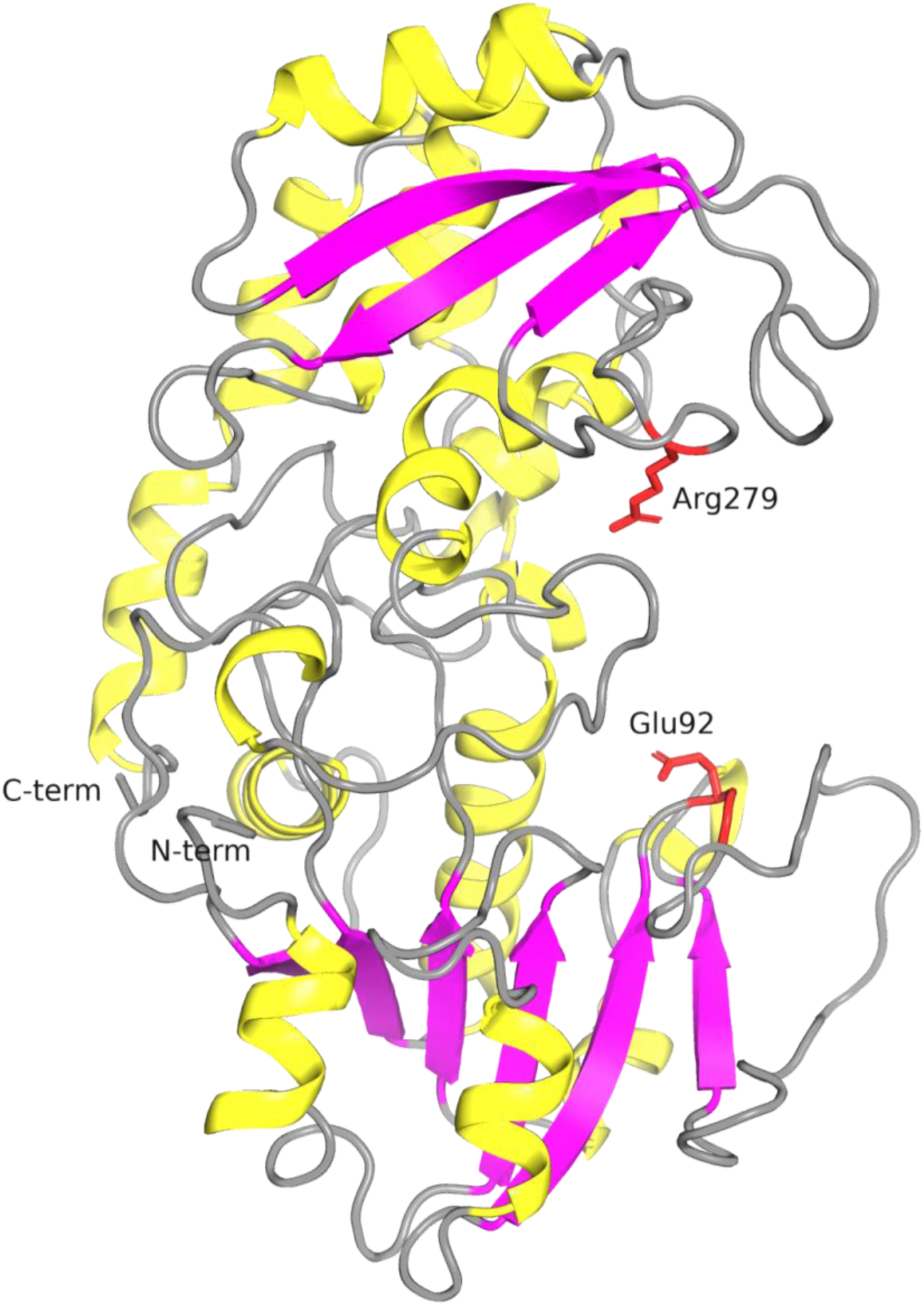
A structure model of *B. subtilis* NamZ depicts a novel glycosidase-fold. A structural model of NamZ, constructed by HHPred, reveals a putative active site located in a cleft within the interface of two sub-domains. The N-terminal catalytic domain contains a Rossmann-fold-like domain (lower part) and contains a conserved glutamate (Glu92) at the tip of the second β-strand. The C-terminal auxiliary domain contains a highly exposed conserved arginine (Arg279).

### Distribution of NamZ and other recycling enzymes

A phylogenetic tree displaying the distribution of recycling associated enzymes, including NamZ, was built as described in Experimental Procedures. Proteins with significant sequence identities to the selected recycling proteins can be found in a wide range of bacterial taxa (Figure 7). Interestingly, NamZ occurs mainly in the phylum of *Bacteriodetes* (in e.g. *Tannerella forsythia*, *Bacteroides fragilis* and *Porphyromonas gingivalis*), and less frequently within Spirochetes (e.g. *Treponema denticola, Leptospira interrogans*), *Bacilli* (*Bacillus subtilis*), *Clostridia* (*Clostridium acetobutylicum*), *Actinobacteria* (*Amycolatopsis mediterranei*, *Brachybacterium faecium*) and *Gammaproteobacteria* (*Yersinia enterocolitica*). Furthermore, the occurrence of NamZ is mostly coupled to the presence of NagZ, whereas, NagZ appears also without NamZ. Also, NamZ and the MurNAc-6P etherase MurQ can be usually found together in the bacterial genomes, since NamZ generates a “precursor substrate” for MurQ. We recently reported on a family of exo-lytic 6-phospho-muramidase that includes MupG of *Staphylococcus aureus*, which specifically hydrolyzes β-1,4-MurNAc-6P entities from the non-reducing end of the disaccharide MurNAc6P-GlcNAc (15). Analysis of the phylogenetic distribution of MupG-like proteins revealed that whenever a putative MurNAc-6P hydrolase is present, NamZ is missing (Figure 7). Considering the complementary functions of the both enzyme families, NamZ and MupG, this observation is reasonable. A characteristic recycling route, the anabolic recycling involving the MurNAc-6P phosphatase MupP, the anomeric MurNAc kinase AmgK, and the MurNAc-α-1P uridylyltransferase MurU has been recently identified in *Pseudomonas sp.* and a range of other Gram-negative bacteria, including *Bacteroides sp.* (35). Analysis of the phylogenetic distribution of recycling enzymes revealed that whenever the anabolic peptidoglycan recycling route is present, NamZ is mostly missing. Thus, NamZ constitutes a PGN degradative enzyme characteristic for a subset of bacterial species, which process PGN by a distinct route that is different from other species lacking NamZ.

**Figure 7.**
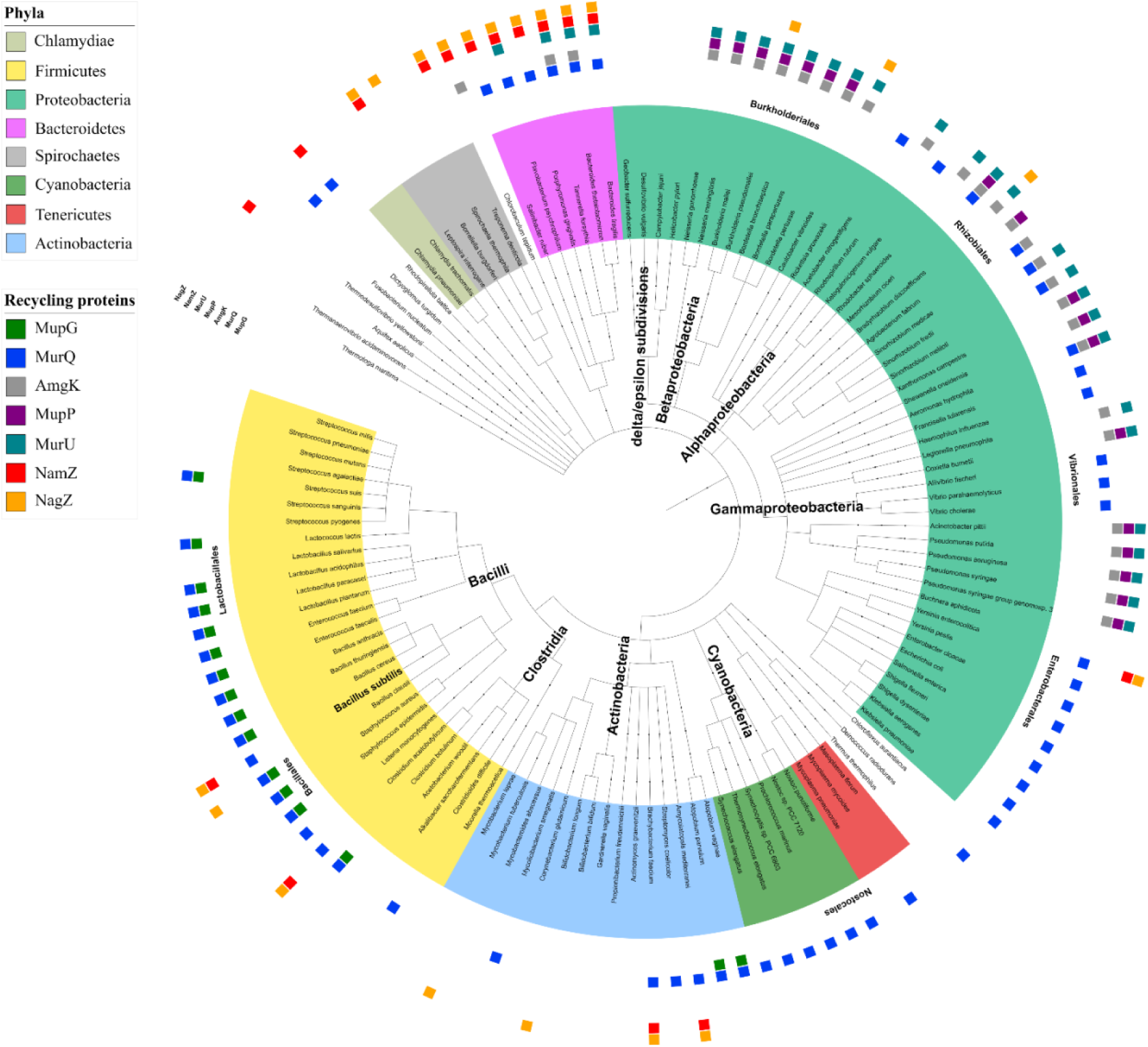
Phylogenetic tree showing the distribution of peptidoglycan recycling associated proteins in representative bacterial taxa. MurQ (blue) is a representative enzyme of the MurQ pathway present in *E. coli* and *B. subtilis* (16,21,22), and MupG (green) is a representative of the modified MurQ pathway identified in *S. aureus* (15). AmgK (grey), MupP (purple) and MurU (teal) are representatives of an alternative, “anabolic” recycling pathway described in *P. putida* (35). NagZ (orange) and NamZ (red) pinpoint the putative presence of autolytic pathways described in *B. subtilis* (16). The tree is built on the taxonomy according to NCBI and MultiGeneBlast searches for these proteins, using the studies by (35) and (50) as reference.

## DISCUSSION

In this study, *ybbC (namZ)* of *B. subtilis* was identified as the first known gene encoding an exo-*N*-acetylmuramidase that specifically cleaves non-reducing terminal MurNAc from synthetic and peptidoglycan-derived natural glycosides. Several lines of evidence indicate that NamZ is identical to the exo-β-*N*-acetylmuramidase that had been partially purified from the supernatant of stationary phase cultures of *B. subtilis* and characterized by Del Rio and colleagues in the 1970s (19,20,31). First, the authors reported that the exo-β-*N*-acetylmuramidase is coexpressed with an exo-β-*N*-acetylglucosaminidase, latter identified as NagZ (16, 20). This is consistent with the location of *ybbC (namZ)* in an operon next to the exo-β-*N*-acetylglucosaminidase *nagZ (ybbD)* and joint regulation of both genes via the transcriptional regulator MurR (YbbH) and catabolite repression (16,21,25). Analysis of growth of Δ*namZ* and Δ*nagZ* mutants and the accumulation of specific turnover fragments, occurring in stationary phase as shown in this study, further supports that NamZ is coexpressed with NagZ. Second, exo-muramidase activity was identified in the growth medium of *B. subtilis* WT but not of Δ*namZ* mutant cells, consistent with the putative signal peptide sequence identified in the N-terminus of the NamZ preprotein and with the previously reported localization and association with cell wall material (20). Third, we showed that NamZ mainly aggregates in solution to form dimers of two 44.77 kDa monomers, consistent with the reported molecular weight of 90 kDa described for the exo-β-*N*-acetylmuramidase by Del Rio *et al.* (20). Fourth, biochemical characteristics and kinetic parameters determined for NamZ further affirm its identity with the exo-β-*N*-acetylmuramidase entity reported by Del Rio *et al*. (19, 20). Del Rio showed the partially purified exo-β-*N*-acetylmuramidase to be maximal active and stable at a pH of 8.0, similarly to our results, that showed an optimum activity of NamZ between pH 6.0 to 8.0 and an optimum stability within the range of pH 6.0 to 10.0. Furthermore, although we used a slightly different artificial substrate with an altered aglycon, the biochemical and kinetic parameters we determined for recombinant NamZ with pNP-MurNAc (*K*_M_ 125 µM; *v*_max_ 1.45 µmol min^-1^ mg^-1^) were very similar to those defined by Del Rio *et al*. using 4MU-MurNAc as substrate (*K*_M_ 190 µM; *v*_max_ 1.5 µmol min^-1^ mg^-1^) (20). The kinetic parameters we determined for MurNAc-GlcNAc, however, differed from those reported by Del Rio *et al.* (20), when the full substrate concentrations range from 0.025 to 6 mM was considered (*K*_M_ 3.6 mM; *v*_max_ 88.64 µmol min^-1^ mg^-1^). Thus, the enzyme appears less affine to the substrate MurNAc-GlcNAc (high *K*_M_), but more active than previous reported (*K*_M_ 650 µM and *v*_max_ 16.29 µmol min^-1^ mg^-1^) (20). However, we observed a biphasic kinetic behaviour with this substrate and if calculating the parameters only for the first saturation, including concentrations up to 0.2 mM MurNAc-GlcNAc (Supporting Figure S8), we obtained kinetic parameters (*K*_M_ 220 µM; *v*_max_ 5.99 µmol min^-1^ mg^-1^) more similar to those reported previously (20). The reason for this discrepancy is not clear but may result from inhibitory or activating compounds that may have been co-purified with the MurNAc-GlcNAc in our or Del Rio’s preparation. We used a similar protocol for substrate preparation, involving peptidoglycan cleavage by the *N*-acetylglucosaminidase Atl^Glc^ of *S. aureus* as well as the subsequent purification of the MurNAc-GlcNAc by liquid chromatography. However, the source of peptidoglycan differed in both preparations, which that was from *B. subtilis* in ours, whereas Del Rio *et al.* used peptidoglycan from *Micrococcus luteus* (20). The major difference is that we removed the peptide stems from the peptidoglycan preparation by treatment with the amidase CwlC from *B. subtilis*, whereas Del Rio only relied on the endogenous amidase present in the *M. luteus* preparation. Although our MurNAc-GlcNAc preparation appears reasonably pure, we cannot exclude that we copurified some cell wall-derived compounds that may affect NamZ kinetics.

The disaccharide MurNAc-β-1,4-GlcNAc likely represents the minimal natural substrate of NamZ. Surprisingly, however, only very little amounts of MurNAc-GlcNAc accumulated in the culture medium of the Δ*namZ* mutant. In the following up of this study, we revealed the reason for this unexpected result to be the fast transport of the disaccharide MurNAc-GlcNAc into the cytoplasm. These results, including the identification of the responsible transporter, will be published elsewhere. Instead, in the growth medium of the Δ*namZ* mutant significant amounts of MurNAc-GlcNAc-anhMurNAc trisaccharide accumulated, about twice the amount that accumulated in the WT strain. Although, the accumulation of the trisaccharide in the WT may indicate that NamZ does not efficiently cleave the anhMurNAc-containing trisaccharide, the MurNAc-GlcNAc-anhMurNAc accumulation product was completely cleavage if recombinant NamZ was added yielding MurNAc and GlcNAc-anhMurNAc products. Thus, cleavage of the non-reducing terminal MurNAc of the trisaccharide by NamZ is, at least in part, responsible for the generation of GlcNAc-anhMurNAc, which accumulated in a huge amount in a Δ*nagZ* mutant. Besides approximately 10-fold lower levels of GlcNAc-MurNAc occur in this mutant. This indicates that, firstly, NagZ is important for the extracellular cleavage of GlcNAc-anhMurNAc and GlcNAc-MurNAc in *Bacillus subtilis*, and, secondly, no uptake of these disaccharides occurs through the cell membrane. The absence of GlcNAc-anhMurNAc and GlcNAc-MurNAc in the culture supernatants of the Δ*namZ* mutant indicates that NamZ is responsible, directly or indirectly, for the generation of these NagZ substrates. Since the amount of MurNAc-GlcNAc-anhMurNAc is rather low, this trisaccharide may not be the main source of GlcNAc-anhMurNAc.

It may be assumed, that primarily long-chain peptidoglycan fragments accumulate in the supernatant of a Δ*namZ* mutant, which are not easily accessible by LC-MS due to their large sizes. Indeed, NamZ as well as NagZ occur associated with cell wall material, and possibly act in concert to degrade peptidoglycan. Our results showed that the exo-lytic *N*-acetylglucosaminidase NagZ and the exo-muramoyl-L-alanine amidase AmiE, along with NamZ can digest intact peptidoglycan, by sequential hydrolysis. However, the rather low amount of thereby released fragments suggest that the decay is not quantitative and likely blocked by structural constraints within the peptidoglycan, such as de-*N*-acetylation. We suppose that the peptidoglycan in *B. subtilis* is autolytically cleaved during vegetative growth by a concerted action of endo-acting as well as disaccharide producing glycosidases (lytic transglycosylases, endo-β-N-acetylmuramidases, and β-N-acetylglucosaminidases), which yield GlcNAc-anhMurNAc, GlcNAc-MurNAc and MurNAc(-peptide)GlcNAc as the final turnover products (Fig. 8). Consistently, we observed an accumulation of GlcNAc-anhMurNAc and GlcNAc-MurNAc in the supernatants of a Δ*nagZ* mutant, and an accumulation of MurNAc-GlcNAc in a Δ*namZ* mutant, which however, as mentioned above, is not found in the culture supernatant since the disaccharide is readily transported into the cells. Candidate peptidoglycan-lytic enzymes, which generate these putative substrates of AmiE, NamZ and NagZ may be the bifunctional lytic transglycosidase/muramidase CwlQ (YjbJ) and the exo-acting *N*-acetylglucosaminidase LytG of *B. subtilis* (36, 37). Altogether, these enzymes yield the monosaccharides GlcNAc and MurNAc, which are taken up and phosphorylated by the PTS transporters MurP and NagP, respectively, and further catabolized intracellularly, in a process known as peptidoglycan turnover and recycling (16, 21). Intracellular MurNAc-6P is converted to GlcNAc-6P by MurQ further enters glycolysis or the cell wall *de novo* synthesis pathway. Consistent with the expression of AmiE, NamZ and NagZ during transition to stationary phase the accumulation of MurNAc-6P occurs during late exponential phase as reported previously (21).

**Figure 8.**
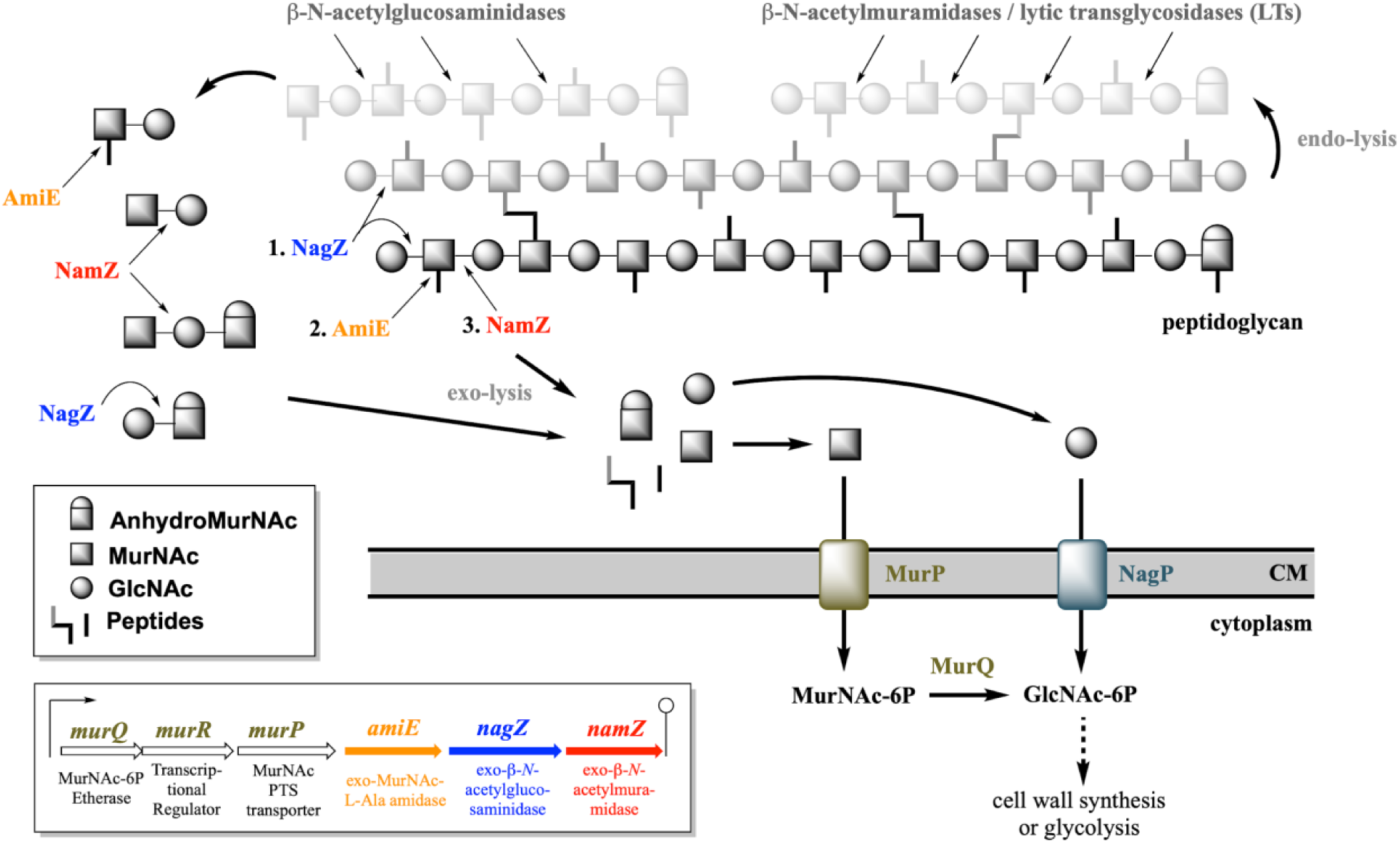
Overview of the roles of NagZ, AmiE, and NamZ of the peptidoglycan recycling/MurNAc catabolic operon in the degradation of peptidoglycan in *B. subtilis*. The exo-lytic peptidoglycan hydrolases NagZ (exo-β-*N*-acetylglucosaminidase), AmiE (exo-*N*-acetylmuramyl-L-alanine amidase) and NamZ (formerly YbbC; exo-β-*N*-acetylmuramidase) cleave-off GlcNAc, peptides, and MurNAc residues, respectively, from the non-reducing terminus of peptidoglycan chains. Furthermore, these enzymes may act on shorter fragments released from the peptidoglycan by the action of endo-lytic and disaccharide-releasing exo-lytic autolysins. Apparently, NamZ primarily cleaves MurNAc-GlcNAc and MurNAc-GlcNAc-anhMurNAc, whease NagZ cleaves GlcNAc-anhMurNAc and GlcNAc-MurNAc. Thereby released monosaccharides, MurNAc and GlcNAc, are recovered by *B. subtilis* via the phosphotransferase system transporters MurP and NagP, yielding in the cytoplasm MurNAc 6-phosphate (MurNAc-6P) and GlcNAc 6-phosphate (GlcNAc-6P), respectively. The MurNAc-6P lactyl ether hydrolase (etherase) MurQ converts MurNAc-6P to GlcNAc-6P in the cytoplasm, which enters pathways leading either to cell wall synthesis or glycolysis.

The exo-*N*-acetylmuramidase NamZ previously belonged to a protein family of so far unknown function (DUF1343) that showed no significant amino acids sequence identity with other glycosidases. A homology model of NamZ that was constructed based on crystal structures of two DUF1343 proteins, BF0379 and BF0371 from *Bacteroides fragilis* NCTC 9343, revealed an atypical glycosidase-fold (Figure 6). A putative active site cleft is located within the interface constituted by two subdomains, a N-terminal Rossmann-fold-like domain and a C-terminal auxiliary α/β domain. Even though NamZ shares higher overall sequence homology with BF0371, the residues around the putative active site of NamZ and BF0379 are more conserved, suggesting that BF0379 and NamZ may share a very similar function. Although the Rossmann-fold-like domain is atypical for glycosidases, the distinguished families 4 and 109 of glycosidases also contain a Rossmann-like-domain fold [URL: https://www.cazypedia.org]. These glycosidases operate by an unusual mechanism that involves an NAD^+^ cofactor required for oxidation/reduction of the hydroxyl group at C3 of the sugar at non-reducing end of a disaccharide to a carbonyl, which renders the C2 proton more acidic and thereby facilitates the elimination of the glycosidic oxygen, i.e. the cleavage of the glycosidic bond (38–41). This unique glycosidase mechanism is, however, incompatible with exo-*N*-acetylmuramidase function, since MurNAc contains a lactyl ether substituent at C3 and not a hydroxyl group. Moreover, we obtained no evidence whatsoever that NamZ function dependents on NAD^+^ and divalent cations, suggesting that the Rossmann-like-fold in DUF1343 enzymes does not involve a NAD^+^ cofactor and a redox-step. Thus, NamZ represents a novel example of a classic fold evolved to adopt new glycoside hydrolase activity. Although some members of glycoside hydrolases (e.g. GH4 and GH109) make use of NAD^+^ as a cofactor that binds to a Rossman-fold domain, to the best of our knowledge, there is no known example of glycoside hydrolase activity arising from a Rossmann-like fold itself. Interestingly, many Rossmann-fold enzymes utilizing nucleoside-based cofactors contain a carboxylate side chain at the tip of the second β-strand (β2-Asp/Glu) for ribose-binding interaction (42), which apparently is conserved in NamZ. Thus, NamZ may represent a novel example of a classic fold evolved to adopt new glycoside hydrolase activity. Exploring the structure/mechanism will be exciting but beyond the scope of the paper and will be addressed in detail in a following up study.

NamZ is a member of a large protein family (pfam PF07075) with presently 2300 predominately bacterial members that have no known function. Thus, NamZ defines a novel family of glycoside hydrolases that have a distinctive structure and possibly mechanism compared to previously known GHs cataloged by the CAZy database. Intriguing, *B. subtilis* is an exception since only very few Firmicutes or Gram-positive bacteria, in general, possess a NamZ protein, which is mostly found in the Baceroidetes phylum (Fig. 7). Importantly, this study fills the gap of the *namZ* (*ybbC*) gene in the peptidoglycan recycling and MurNAc catabolic operon and provides insights into the role of NagZ, AmiE, and NamZ in peptidoglycan turnover by characterizing its NamZ glycosidase protein product and assigning its unique exo-muramidase function.

## EXPERIMENTAL PROCEDURES

### Chemicals, media, enzymes, and oligonucleotides

Chemicals were obtained from AppliChem GmbH (Darmstadt, Germany), Bachem (Bubendorf, Switzerland), Carbosynth Ltd. (Berkshire, UK) Serva Electrophoresis GmbH (Heidelberg, Germany), Sigma-Aldrich (St. Louis, USA), Thermo Fisher Scientific (Waltham, Massachusetts, USA), Carl Roth GmbH + Co. KG (Karlsruhe, Germany) and VWR (Radnor, Pennsylvania, USA). Antibiotics were received from Sigma-Aldrich (St. Louis, USA) and lysogeny broth medium (LB broth Lennox) was from Carl Roth GmbH + Co. KG (Karlsruhe, Germany). DNA polymerases and enzymes for DNA restriction and cloning were from Genaxxon bioscience GmbH (Ulm, Germany), New England Biolabs (Ipswich, USA) and Thermo Fisher Scientific (Waltham, Massachusetts, USA). The Gene JET Genomic DNA Purification Kit, Plasmid Miniprep Kit, PCR Purification Kit, and GeneRuler 1-kb marker were also from Thermo Fisher Scientific (Waltham, Massachusetts, USA). Oligonucleotides and sequencing results were received from MWG Eurofins (Ebersberg, Germany).

### Bacterial strains and growth conditions

The bacterial strains and plasmids used in this study are listed in Table 1. For the generation of markerless *B. subtilis* 168 Δ*nagZ* and Δ*namZ* mutants, the loop-out plasmid pDR244 was used to excise the erythromycin resistance cassettes flanked by *loxP* sites in strains *B. subtilis* 168-Δ*nagZ::erm* and *B. subtilis* 168-Δ*namZ::erm,* received from the *Bacillus* Genetic Stock Center (Columbus, Ohio, USA), following a published protocol for constructing markerless deletions. In brief, the preparation of naturally competent cells and transformation of pDR244, carrying the *cre* recombinase gene, was accomplished using MGE medium (43), followed by selection on spectinomycin (100 µg/ml) and incubation at 30°C overnight. Transformant colonies were shifted to incubation at 42°C overnight on LB without added antibiotic. Growing colonies were streaked out on LB, spectinomycin (100 µg/ml) and erythromycin (5 µg/ml) to verify *cre*-dependent loss of the erythromycin resistance cassette and plasmid curing. Colonies growing only on LB were confirmed for the loss of the erythromycin resistance cassette by PCR using *B. subtilis* Δ*nagZ* and *B. subtilis* Δ*namZ* chromosomal DNA and primers flanking the regions of interest (nagZ_500_fw and nagZ_500_rev for Δ*nagZ* as well as namZ_500_fw and namZ_500_rev for Δ*namZ*; Table S2) as well as DNA sequencing of the amplicons. All strains were cultured under aerobic conditions at 37°C in LB under continuous shaking at 140 rpm. Antibiotics were used when appropriate at the following concentrations (50 µg/ml kanamycin or 100 µg/ml ampicillin).

### Analysis of supernatants of *B. subtilis* WT, Δ*namZ* and Δ*nagZ cells*

*B. subtilis* wild type, Δ*namZ* and Δ*nagZ* cells were grown for 20 h under constant shaking. Afterwards, 1 ml cell cultures were harvested and centrifuged at 12000 x g000 x g for 10 min at room temperature and 100 µl of the cell-free supernatant was added to 900 µl of ice-cold acetone. Samples were mixed by inverting the tubes three times. Following precipitation, the samples were centrifuged at 12000 x g for 10 min. The supernatants were dried at 55°C (SpeedVac, Christ, RVC2-18, Osterode am Harz, Germany) and the resulting pellet was resuspended in 50 µl milliQ. Finally, 3 µl of the bacterial supernatants were analyzed by HPLC-MS.

### HPLC-MS analysis of supernatants, pNP-MurNAc, MurNAc-GlcNAc and digested peptidoglycan

Sample analysis of digested peptidoglycan samples was accomplished using an electrospray ionisation-time of flight (ESI-TOF) mass spectrometer (MicrOTOF II, Bruker Daltonics), operated in positive or negative ion mode, connected to an high performance liquid chromatography (HPLC) system (UltiMate 3000, Dionex). For HPLC-MS analysis samples were injected (3 µl for MurNAc-GlcNAc samples, MurNAc standard and for supernatants, 5 µl for pNP-MurNAc and digested peptidoglycan samples) onto a Gemini C18 column (150 x 4.6 mm, 5 µm, 110 Å, Phenomenex) and separated at 37°C with a flow rate of 0.2 ml/min in accordance with a previously described program (35). The mass spectra of analyzed samples were displayed as extracted ion chromatograms (EIC) in DataAnalysis program (Bruker) and were presented in Prism 8 (GraphPad). The relative amounts of MurNAc from MurNAc standards and from release of MurNAc-GlcNAc substrate in enzyme kinetic studies were determined by calculating the area under the curve (AUC) of the corresponding EIC spectra for MurNAc.

### Plasmid constructions for heterologous expression of *namZ*, *amiE, cwlC*, and *atl^Glc^*

Genomic DNA of *B. subtilis* 168 and *S. aureus* USA300 JE2, serving as PCR-templates, was prepared using GeneJET Genomic DNA Purification Kits. The genes *namZ, amiE, and cwlC* from *B. subtilis* 168 and a*tl^Glc^* from *S. aureus* USA300 JE2 were amplified by PCR using Phusion DNA polymerase using the primer pairs ybbC_FW/ybbC_RV, RK-24/RK-20, AmiE_Bs_for/AmiE_Bs_rev and 69_Atl_gluc_Fw/70_Atl_gluc_Rev, respectively (listed in Table S2). In all cases the natural signal sequence was removed, allowing cytoplasmic expression in *E. coli*. Cloning and expression of *amiE* of *B subtilis* along with *nagZ* has been achieved before (16). However, we decided to reclone *amiE* in a pET vector system for an easy, collective expression of all required enzymes in this study. PCR products and vectors were digested with the corresponding restriction enzymes, using *Nde*I and *Xho*I for *namZ* with pET29b, and *Nco*I and *Xho*I for *cwlC, atl^Glc^* with pET28a, as well as for *amiE* with pET22b. Accordingly, using T4 DNA ligase *namZ* was ligated into a pET29b expression vector, *cwlC* and *atl^Glc^* into pET28a expression vectors, and *amiE* into a pET22b expression vector (all vectors were from Novagen, Darmstadt, Germany). Thereby all recombinant proteins were provided with a C-terminal His_6_-tag. Chemically competent *E. coli* DH5α cells were transformed with the ligation reaction mixtures. Recombinant plasmids pET29b-*namZ*, pET28a-*clwC*, pET28a-*atl^Glc^* and pET22b-*amiE* were isolated from kanamycin- or ampicillin-resistant *E. coli* cells and DNA sequences were verified by sequencing. Chemically competent *E. coli* BL21(DE3) cells were then transformed with pET29b-*namZ*, pET28a-*clwC*, pET28a-*atl^Glc^* and pET22b-*amiE* and used to heterologously express the recombinant proteins under the control of the T7 promotor.

### Expression and purification of recombinant NamZ, NagZ, AmiE, CwlC, and Atl^Glc^

For the overexpression of NamZ and AmiE 2L LB medium and for overexpression of CwlC and Atl^Glc^ 1L LB medium, all supplemented with 50 µg/ml kanamycin or 100 µg/ml ampicillin (for pET22b-amiE) were inoculated with overnight cultures of *E. coli* BL21(DE3) harboring the expression plasmids pET29b-*namZ*, pET28a-*cwlC*, pET28a-*atl^Glc^* and pET22b-*amiE*. The cells were cultivated at 37°C under continuous shaking. Expression of the recombinant proteins was induced at log phase (at OD_600_ 0.7) by addition of IPTG at a final concentration of 1 mM. After induction the cells harboring pET29b-*namZ*, pET28a-*cwlC* and pET28a-*atl^Glc^* were grown further for 3 h at 37°C, and the cells harboring pET22b-*amiE* were grown overnight at 18°C, all under continuous shaking. Cells were harvested by centrifugation at 4000 x g for 30 min at 4°C. Cell pellets were resuspended with 20 ml of sodium phosphate buffer A (20 mM Na_2_HPO_4_ * 2 H_2_O, 500 mM NaCl, pH 7.5) each and disrupted by using French Press (Sim Aminco Spectronic Instruments, Inc. Rochester, New York, USA) three times at 1000 psi. Cell debris and unbroken cells were removed by centrifugation at 38000 x g for 60 min at 4 °C. Purification of the C-terminally His_6_-tagged proteins was performed by Ni^2+^ affinity chromatography. Therefore the obtained supernatants were filtered through 0.2 μm filters (Sarstedt, Nümbrecht, Germany) and loaded on 1 ml His-Trap columns (GE Healthcare, Freiburg, Germany), pre-equilibrated with ten column volumes of each ddH_2_O and sodium phosphate buffer A (20 mM Na_2_HPO_4_ * 2 H_2_O, 500 mM NaCl, pH 7.5), using a protein purification system (ÄKTA purifier, GE Healthcare). Elution of the proteins was achieved by using a linear gradient of imidazole from 0 to 500 mM with sodium phosphate buffer B (20 mM Na_2_HPO_4_ * 2 H_2_O, 500 mM NaCl, 500 mM imidazole, pH 7.5). Purity of the enzymes was analyzed by SDS-PAGE (12% polyacrylamide) stained with Coomassie Brilliant Blue G250. Peak fractions containing desired proteins were pooled and further purified by size exclusion chromatography (HiLoad 16/60 Superdex 200 column, GE Healthcare) using sodium phosphate buffer A as eluent. Peak fractions were analyzed for pure proteins by SDS-PAGE (12% polyacrylamide) and fractions containing pure enzymes were pooled. Protein concentrations were determined using the extinction coefficient at 280 nm (36330 M^-1^ cm^-1^ for NamZ, 15930 M^-1^ cm^-1^ for CwlC, 97180 M^-1^ cm^-1^ for Atl^Glc^ and 50770 M^-1^ cm^-1^ for AmiE, ExPASy, ProtParam tool) and measured in a 1 ml quartz cuvette (Hellma, Müllheim, Germany) using a SpectraMax M2 spectrophotometer (Molecular Devices, Biberach, Germany). NagZ was purified according to a previously published protocol (16).

### Size exclusion chromatography and multi angle light-scattering (SEC-MALS)

Analytical size exclusion chromatography (SEC) coupled to a multi angle light scattering apparatus (MALS, Wyatt Technology Corp., Dernbach, German) was used to determine the apparent molecular masses of NamZ protein as described previously in (44). The experiments were done on a micro-Äkta chromatography system (GE Healthcare) connected to SEC Superose 6 Increase column (10/300 GL, GE Healthcare) at room temperature with a flow rate of 0.5 ml/min in sodium phosphate buffer A (20 mM Na_2_HPO_4_ * 2 H_2_O, 500 mM NaCl, pH 7.5). MALS analysis was achieved by connecting micro-Äkta to a downstream MALS apparatus to which a refractometer (Optilab T-rEX, Wyatt Technology) and a miniDawn Treos system (Wyatt) were connected. The UV signals were plotted against the elution volume and the molar masses were calculated from the light scattering data using ASTRA 7 software (Wyatt) and the mass distribution was plotted over the eluted peaks.

### Determination of the substrate specificity of recombinant NamZ with chromogenic substrates

NamZ activity was shown by cleavage of the chromogenic substrate para-nitrophenyl 2-acetamido-3-O-(D-1-carboxyethyl)-2-deoxy-β-D-glucopyranoside (pNP-MurNAc). Specificity of NamZ was further examined by testing para-nitrophenyl 2-acetamido-2-deoxy-β-D-glucopyranoside (pNP-GlcNAc) (Sigma-Aldrich, St. Louis, USA) and o-nitrophenyl β-D-galactopyranoside (oNP-Galactose) (Carl Roth GmbH + Co. KG, Karlsruhe, Germany) as substrates of NamZ. As positive control enzymes, NagZ (exo-β-*N*-acetylglucosaminidase of *B. subtilis*, purified according to (16), and LacZ (β-D-galactosidase of *E. coli* (Hoffmann-La Roche, Basel, Switzerland) were used. In the test assay for NamZ specificity, a 100 µl reaction volume, containing chromogenic substrates (100 µM each) as listed above in 0.2 M sodium phosphate buffer (pH 8.0) was incubated for 30 min at 37 °C with purified enzymes (5 µg NamZ (0.7 µM), 5 µg NagZ (0.48 µM) and 10 µl LacZ (1500 U)). The reactions were initiated by addition of 10 µl of enzyme and stopped after 30 min by the addition of 100 µl borate buffer (0.5 M, pH 10.0). Specific release of the nitrophenol groups could be visualized by a yellow color and absorption was additionally measured at 405 nm. The assay was also conducted with pNP-isoMurNAc, which however showed no yellow color release with none of the enzymes, including NamZ.

### Determination of temperature and pH stability and optima of NamZ

Temperature stability of NamZ was determined by pre-incubating NamZ at different temperatures (i.e., 4, 20, 37, 45, 55 and 60 °C) for 30 min, and subsequently measuring the kinetics of the NamZ reaction under standard conditions (200 µM pNP-MurNAc in 200 mM sodium phosphate buffer, pH 7.5 at 30 °C). Therefore, 10 µl each of the pre-incubated enzyme (500 ng) was added to 90 µl reaction mixtures containing 200 µM pNP-MurNAc in 200 mM sodium phosphate buffer (pH 7.5) and the reaction at 37 °C was followed by measuring the absorption at 405 nm in a SpectraMax M2 microplate reader (Molecular Devices, Biberach, Germany). Determination of temperature optimum of NamZ was achieved, by adding 10 µl of fresh enzyme (500 ng, 105 nM) to 90 µl reaction mixtures containing 200 µM pNP-MurNAc in 200 mM sodium phosphate buffer (pH 7.5) directly following the reaction at different temperatures (i.e., 4, 20, 37, 45, 55, 60 °C) by measuring the absorption at 405 nm.

Determination of pH stability and pH optimum of NamZ was achieved, by applying buffers in the pH range of 2.0 to 10.0, i.e., Clark and Lubs buffer (pH 2.0), citric acid-sodium phosphate buffer (pH 3.0 to 7.0), hydrochloric acid-Tris buffer (pH 8.0) and glycine-sodium hydroxide buffer (pH 9.0 to 10.0). For determination of pH stability, NamZ was diluted in appropriate buffers ranging from pH 2.0 to 10.0 at a final concentration of 50 ng/µl. The enzyme was pre-incubated for 30 min at ambient temperature (22 °C). After incubation the mixtures were buffer exchanged in centrifugal filter units (Amicon Ultra, 0.5 ml, 10 K, Merck Millipore, Darmstadt, Germany) washed three times with 50 µl of sodium phosphate buffer (20 mM Na_2_HPO_4_, 500 mM NaCl, pH 7.5). After washing, the volume was brought up to 50 µl with 200 mM sodium phosphate buffer (pH 7.5) to maintain a final enzyme concentration of 50 ng/µl. The reaction assay was initiated by adding 10 µl of pre-incubated enzyme (500 ng, 105 nM) to a 90 µl reaction mixture under standard condition (as described above). For determination of pH optimum 90 µl reaction mixtures were prepared containing 200 µM pNP-MurNAc in buffers of a particular pH in the range of 2.0 to 10.0. The reaction was initiated by adding 10 µl enzyme (500 ng, 105 nM) followed by incubation for 60 min at 37 °C; the reaction was stopped with borate buffer (250 mM disodium tetraborate, 1 M NaOH, pH 10.8). Samples were analyzed as described above.

### Preparation of peptidoglycan

Peptidoglycan was required on the one hand to generate MurNAc-GlcNAc as natural substrate for the biochemical characterization of NamZ and on the other hand to generate polymeric substrate for the sequential digest using NagZ, AmiE and NamZ. Peptidoglycan of *B. subtilis* was prepared as previously described with slight modifications (45). For the production of MurNAc-GlcNAc, two 5 L liter Erlenmeyer flasks each containing 1 L LB medium were inoculated with *B. subtilis* cells to yield an initial OD_600_ of 0.1. The cells were cultured at 37 °C with continuous shaking at 120 rpm and harvested at an OD_600_ of 1.8 by centrifugation at 5000 x g for 15 min at 4 °C. For the preparation of the peptidoglycan substrate for the sequential digest using NagZ, AmiE, and NamZ, exponential phase cells were harvested at an OD_600_ of 1.0 by centrifugation at 5000 x g for 15 min at ambient temperature. For the preparation of MurNAc-GlcNAc, the cells were washed with 25 ml PBS (137 mM NaCl, 2.7 mM KCl, 10 mM Na_2_HPO_4_, 1.8 mM KH_2_PO_4_) and the resulting pellets were stored at −20 °C. Frozen cells were suspended in 20 ml phosphate buffer (25 mM, pH 6.0) and added dropwise to a boiling solution (20 ml) of 8% SDS solution in phosphate buffer (25 mM, pH 6.0). After boiling under constant stirring for 30 min the samples were cooled down to ambient temperature and centrifuged at 50’000 x g for 30 min at 40 °C. The pellet was washed with a phosphate buffer (25 mM, pH 6.0) until the SDS in the sample was completely removed. The removal of SDS was proven with a methylene blue assay (46). The SDS-free pellet was suspended in 10 ml phosphate buffer (25 mM, pH 6.0) and incubated with 100 µg/ml α-amylase (from *Bacillus sp.*, Sigma-Aldrich, St. Louis, USA) for 1 h at 37 °C. Afterwards, 200 µg/ml pronase (from *Steptomyces griseus*, Calbiochem, Merck KGaA, Darmstadt, Germany) were added and the sample was incubated overnight at 37 °C. At the next day, the sample was centrifuged at 16’500 x g for 10 min and the pellet was incubated with 5 ml of 1 M HCl for 4 h at 37 °C to remove the wall teichoic acids. After the HCl treatment the pellet was washed with ddH_2_O (16’500 x g, 10 min) until a pH of 5 to 6 was reached. The pellet was suspended in 20 ml phosphate buffer (25 mM, pH 6.0) and added dropwise into 20 ml of boiling phosphate buffer (25 mM, pH 6.0) and SDS (4%). Removal of SDS in the sample was performed as previously described.

For the peptidoglycan preparation for the sequential digests, the fresh cells were resuspended in 15 ml of Tris-HCl buffer (50 mM, pH 7.5) containing 20 µl proteinase K (20 mg/ml, NGS grade, 7BioScience GmbH, Hartheim, Germany) and added dropwise into 15 ml of boiling Tris-HCl buffer (50 mM, pH 7.5). After boiling for 60 min the sample was cooled down to room temperature and centrifuged at 2900 x g for 15 min at room temperature. The pellet was stored at −20 °C until the next day. Frozen pellet was resuspended in 6 ml Tris-HCl buffer containing MgSO_4_ (50 mM, 10 mM MgSO_4_, pH 7.5) and incubated together with 600 µg α-amylase (from *Bacillus sp.*, Sigma-Aldrich, St. Louis, USA), 10 µl RNase A (10 mg/ml, Thermo Fisher Scientific, Waltham, Massachusetts, USA) and 10 µl DNase I (5 U/µl, Thermo Fisher Scientific, Waltham, Massachusetts, USA) for 2 h at 37 °C under continuous shaking. After incubation, 20 µl proteinase K (20 mg/ml, NGS grade, 7Bioscience GmbH, Hartheim, Germany) was added to the suspension, and immediately added dropwise into boiling SDS solution (end-concentration of SDS was 2%), followed by boiling for 60 min. To remove the residual SDS, the sample was centrifuged in ultracentrifugation bottles (Beckman Coulter, Pasadena, California, USA) at 104500 x g at 40 °C for 30 min. The pellet was resuspended in pre-warmed ddH_2_O (about 60 °C) and centrifuged at least 10 times in an ultracentrifuge (Beckman Coulter, Pasadena, California, USA), until residual SDS was completely removed. To verify the removal of SDS from peptidoglycan samples, a methylene blue assay was performed (46). To remove wall teichoic acids, the peptidoglycan pellet was resuspended in 2 ml of 48 % hydrofluoric acid (Sigma-Aldrich, St. Louis, USA) and incubated for 48 h at 4 °C under vigorous shaking. Afterwards, the sample was washed several times with phosphate buffer (100 mM, pH 7.0, 20000 x g, 5 min) and the peptidoglycan pellet was neutralized with ddH_2_O until the pH of the supernatant was above 6.5. Finally, the purified peptidoglycan was dried (SpeedVac, Christ, RVC2-18, Osterode am Harz, Germany) and weighed.

### Purification of MurNAc-GlcNAc

To generate MurNAc-GlcNAc, 900 mg peptidoglycan were incubated with CwlC and Atl^Glc^ (each 1 µM) in 90 ml sodium phosphate buffer (100 mM, pH 8.0) overnight at 37 °C under continuous shaking. Afterwards, the reaction mixture was heated to 95 °C for 25 min. 90 µl of the heat-treated samples were loaded on a semi-preparative Gemini C18 column (250 x 10 mm, 110 Å, 5 μm, Phenomenex) and separated via reversed-phase HPLC (VWR Hitachi Chromaster, Germany) at 20 °C with a flow rate of 1.5 ml/min in accordance with a previously described program (35) with slight modifications, starting with 10 min of washing with 100% buffer A (0.1% formic acid, 0.05% ammonium formate in water), afterwards injection of the sample and another 5 min washing step with 100% buffer A. A linear gradient to 40% buffer B (acetonitrile) over 30 min was followed by a 5 min hold at 40% buffer B. Re-equilibration within a 1 min step to 100% buffer A was followed by a 10 min wash step at 100% buffer A. Fractions were collected in 1.5 ml volumes, representing 1 minute of flow each. Collected fractions containing MurNAc-GlcNAc were lyophilized, suspended in 3 ml ddH_2_O, aliquoted in tubes and dried at 40 °C using SpeedVac (Christ, RVC2-18, Osterode am Harz, Germany).

Specificity of NamZ was determined using natural cell wall disaccharides MurNAc-GlcNAc (as prepared above) and GlcNAc-MurNAc (Carbosynth Ltd., Berkshire, UK). For each substrate, two 25 µl reaction mixtures were prepared containing either 100 µM MurNAc-GlcNAc or 100 µM GlcNAc-MurNAc in 0.2 M sodium phosphate buffer (pH 7.0). The reactions were initiated by adding 500 ng purified NamZ (420 nM). In addition, NagZ (500 ng, 280 nM) was added as a control to the reaction mixtures. As a negative control, reaction mixtures without added enzymes were incubated. The reaction mixtures were incubated at 37 °C for 30 minutes and stopped with a buffer containing 1% formic acid and 0.5% ammonium formate in water (pH 3.2). The reaction mixture was transferred into vials and analyzed directly by HPLC-MS under conditions as described above.

### Determination of enzyme kinetic parameters for pNP-MurNAc and MurNAc-GlcNAc substrates

Determination of kinetic parameters of NamZ were achieved by using both, the chemically synthesized substrate pNP-MurNAc, and the purified substrate MurNAc-GlcNAc. Before starting the kinetic experiments, the purity of pNP-MurNAc was checked by HPLC-MS analysis. For pNP-MurNAc, 100 µM substrate in phosphate buffer (0.2 M, pH 7.0) were analyzed and compared to the same reaction mixture containing additionally 500 ng NamZ (105 nM). Both mixtures were incubated at 37 °C for 30 min and subsequently analyzed using HPLC-MS. Since we discovered that the pNP-MurNAc stock in MeOH formed traces of methyl ester, which could not serve as a substrate for NamZ, we chose incubation under basic conditions to fully hydrolyse the ester traces of pNP-MurNAc. Therefore, 1 mM pNP-MurNAc were pre-incubated in phosphate buffer (20 mM, pH 8.0) at 37 °C for 30 min. Afterwards, 100 µM pre-incubated pNP-MurNAc in phosphate buffer (0.2 M, pH 7.0) were analyzed and compared to the same reaction mixture containing additionally 500 ng NamZ (420 nM). Both mixtures were, again, incubated at 37 °C for 30 min and subsequently analyzed using HPLC-MS.

To ascertain the kinetic parameters for pNP-MurNAc, different concentrations of pNP-MurNAc in 0.2 M phosphate buffer (pH 8.0) were used, after pre-incubation in phosphate buffer as described above. In a 96-well plate (Greiner, Frickenhausen, Germany), a 90 µl reaction mixture containing pNP-MurNAc ranging from 13 to 860 µM in phosphate buffer (0.2 M, pH 8.0) was incubated at 37 °C. The reaction was started by addition of 10 µl enzyme (500 ng, 105 nM). The reaction was continuously monitored at 37 °C for 30 min at 405 nm with a Spark 10 M microplate reader (Tecan, Männedorf, Switzerland). To determine the amount of released 4-nitrophenol by the action of NamZ on pNP-MurNAc, a 4-nitrophenol standard (Sigma Aldrich, St. Louis, USA) dissolved in phosphate buffer (0.2 M, pH 8.0) with concentrations from 13 to 430 µM was also prepared for measurement at 405 nm with the same microplate reader as mentioned above. The experimental data of the enzyme kinetics were evaluated by nonlinear regression of the reaction curve using the program Prism 6 (GraphPad).

To determine the kinetic parameters for MurNAc-GlcNAc, different concentrations of MurNAc-GlcNAc in 0.2 M phosphate buffer (pH 7.0) were used. In eppendorf tubes (Sarstedt, Nürnbrecht, Germany), a 18 µl reaction mixture containing MurNAc-GlcNAc ranging from 0.025 to 6 mM in phosphate buffer (0.2 M, pH 7.0) was incubated at 37 °C. The reaction was started by adding 2 µl enzyme (10 ng, 10.5 nM). The reaction mixture was incubated at 37 °C for 5 minutes and stopped with 20 µl of stopping buffer containing 1% formic acid and 0.5% ammonium formate in water (pH 3.2). The reaction mixture was transferred into vials and analyzed by HPLC-MS. To determine the amount of released MurNAc by the action of NamZ on MurNAc-GlcNAc, a MurNAc standard with concentrations from 0.0063 to 0.8 mM was also prepared for HPLC-MS analysis. To ensure the same ionization conditions for the standard and the reaction mixture, the standard was dissolved in an equal mixture of 0.2 M phosphate buffer (pH 7.0) and the stopping buffer.

### Sequential digest with NagZ, AmiE and NamZ

To perform the sequential digest, 2.5 mg of purified *B. subtilis* peptidoglycan were dissolved in 100 µl phosphate buffer (0.2 M, pH 7.0). To the dissolved peptidoglycan a 50 µl mixture containing 2.5 µM NagZ (26.7 µg), 2.5 µM AmiE (17.9 µg) and 2.5 µM NamZ (19.5 µg) in phosphate buffer (0.2 M, pH 7.0) was added, resulting in a total volume of 150 µl. The digest was incubated overnight at 37 °C under continuous shaking. After incubation the reaction was stopped by heating at 95 °C for 30 min. To separate cell wall monosaccharides and peptides from undigested peptidoglycan, the reaction mixture was centrifuged at 12100 x g for 15 min and the supernatant was dried at 37 °C (SpeedVac Christ, RVC2-18, Osterode am Harz, Germany). Afterwards the pellets were dissolved in 50 µl ddH_2_O and transferred into vials to be analyzed using HPLC-MS.

### Taxonomic distribution of NamZ and other recycling enzymes

A taxonomic tree was constructed that depicts the distribution of homologs of the recycling proteins NamZ and NagZ of *B. subtilis* 168, as well as other recycling proteins, MurQ of *E. coli* K12, MupG of *S. aureus* USA300, and AmgK, MupP, and MurU of *P. putida* KT2440. The occurrence of the homologs was investigated in organisms included on the representative NCBI RefSeq genomes. The protein sequences for NagZ (P40406) and NamZ (P40407), as well as MurQ (P76535), MupG (A0A0H2XHV5), AmgK (Q88QT3), MupP (Q88M11), and MurU (Q88QT2), were obtained from UniProt and BLAST searches were analyzed using the MultiGeneBlast application (47). MultiGeneBlast allows homology searches with more than one input sequence and also displays the localization of genes coding for possible homologues in the genome. The application was run in “architecture search” mode to find the genes coding for possible homologues within a distance of 20 kb, regardless of their co-localization or order of occurrence on the genome. Furthermore, thresholds were set for a minimum sequence coverage of 50% and a minimum identity of 20%, leaving the other settings at the default values. Matches found were annotated and visualized in a tree constructed according to the NCBI taxonomy using the iTOL v3 web service (48).

## Acknowledgements

C.M. acknowledges funding by the ministry of science, research and art of the state Baden-Württemberg (programme glycobiology/glycobiotechnology) and by the Deutsche Forschungsgemeinschaft (DFG, German Research Foundation), grants SFB766, Project-ID 398967434 – TRR 261 and Project-ID 174858087 – GRK1708. This work was further supported by the DFG-funded Cluster of Excellence EXC 2124 Controlling Microbes to Fight Infections. We thank Amanda Duckworth for the construction of plasmid pET29b-namZ and Libera lo Presti for manuscript editing.

## Author contributions

CM conceptualization; MM, MC, IH, QX and MB data curation; MM, MC, IH, RMK, TT and MB formal analysis; CM funding acquisition; MM and CM investigation; MM, MC, IH, RMK, KAS, and CM methodology; CM project administration; MM, MC, RMK, TT, KB, AE, and QX resources; MB, AT and CM supervision; MM, MC, MB and AT validation; MM, MC, MB and CM visualization; MM and CM writing-original draft; MB, AT and CM writing-review and editing.

## Footnotes

The abbreviations used are: MurNAc, *N*-acetylmuramic acid; GlcNAc, *N*-acetylglucosamine; pNP-GlcNAc, para-nitrophenyl 2-acetamido-2-deoxy-β-D-glucopyranoside; pNP-MurNAc, para-nitrophenyl 2-acetamido-3-O-(D-1-carboxyethyl)-2-deoxy-β-D-glucopyranoside; EIC, extracted ion chromatogram; AUC, area under curve; BPC, base peak chromatogram

## SUPPORTING INFORMATION

### Supporting Experimental Procedures

#### Synthesis of pNP-MurNAc and pNP-isoMurNAc

The synthesis of the chromogenic substrate para-nitrophenyl-2-acetamido-3-O-(D-1-carboxyethyl)-2-deoxy-β-D-glucopyranoside (pNP-MurNAc) and its stereoisomer para-nitrophenyl-2-acetamido-3-O-(L-1-carboxyethyl)-2-deoxy-β-D-glucopyranoside (pNP-isoMurNAc) was carried out according to modified literature procedures (1).

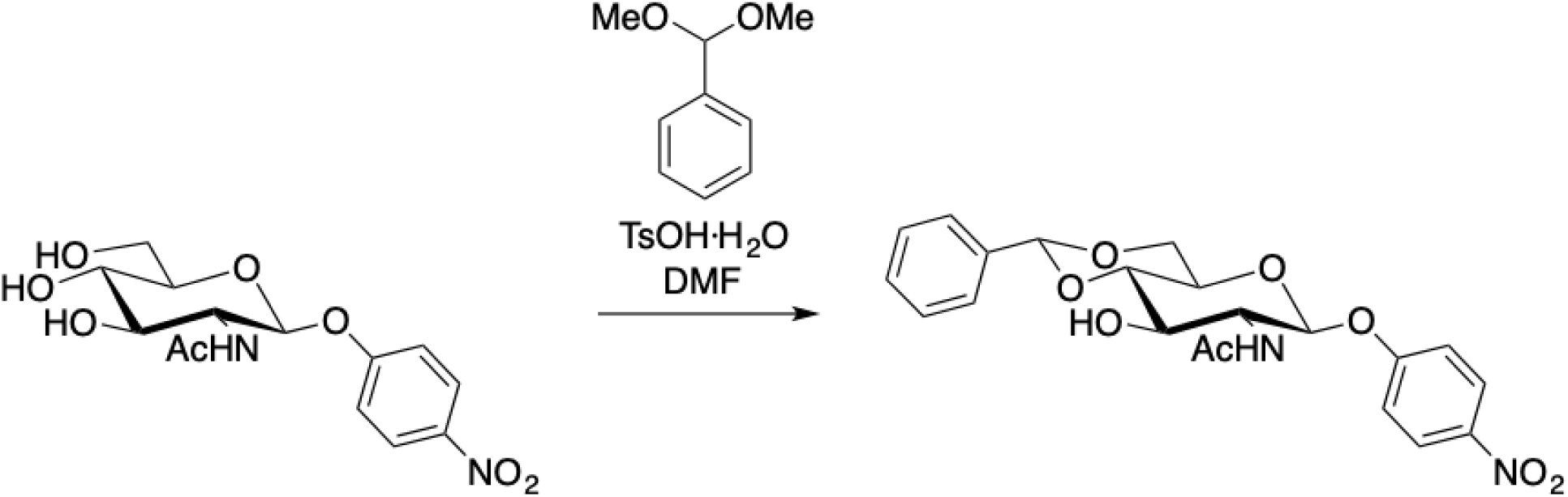

#### p-Nitrophenyl 4,6-benzylidine-GlcNAc

According to the literature procedure, (2) p-nitrophenyl GlcNAc (220 mg, 0.64 mmol) in DMF (3 mL) was treated with benzaldehyde dimethyl acetal (120 µl, 0.78 mmol) and TsOH·H2O (9 mg). The suspension was rotated on a rotary evaporator (50 °C, 40 mbar) for 2.5 hours. The clear solution was treated with NaHCO3 (saturated aqueous solution, 40 ml) and stirred at room temperature for 30 minutes. The precipitate was collected by filtration and washed with water and ether to yield the title compound (214 mg, 77%) as a colourless solid. ^1^H NMR (500 MHz, DMSO-*d*_6_) δ 8.20 (d, *J* = 9.3 Hz, 2H), 8.02 (d, *J* = 8.8 Hz, 1H), 7.50 – 7.36 (m, 3H), 7.23 (d, *J* = 9.2 Hz, 2H), 5.64 (s, 1H), 5.39 (d, *J* = 8.4 Hz, 1H), 4.29 – 4.19 (m, 1H), 3.88 – 3.79 (m, 1H), 3.79 – 3.68 (m, 3H), 3.56 (t, *J* = 8.9 Hz, 1H), 1.81 (s, 3H). ^13^C NMR (126 MHz, DMSO-*d*_6_) δ 169.52, 161.90, 142.05, 137.66, 128.98, 128.12, 126.39, 125.86, 116.64, 100.77, 98.43, 80.80, 70.16, 67.67, 66.19, 55.89, 23.06. Analytical data in agreement with the literature (3).

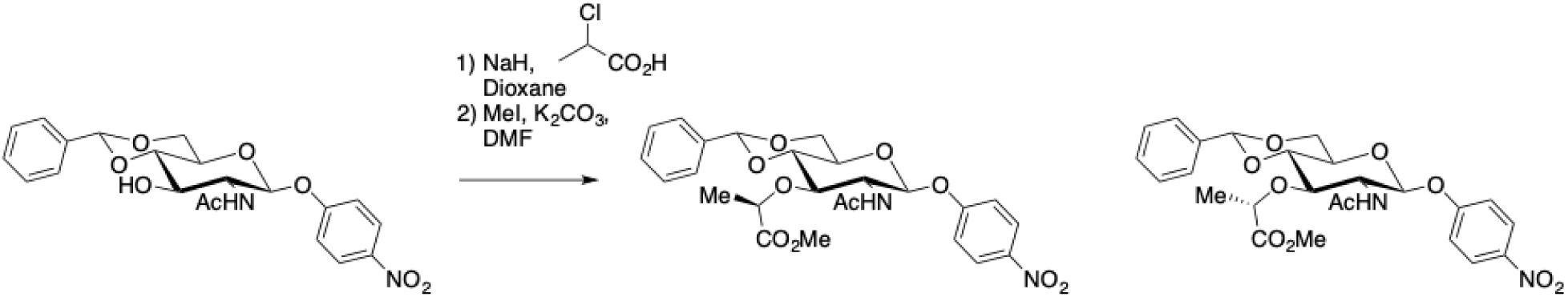

#### p-Nitrophenyl 3-O-(D-1-methoxycarbonylethyl)-4,6-benzylidine-GlcNAc

p-Nitrophenyl 4,6-benzylidene-GlcNAc (171 mg, 0.4 mmol) was taken up in toluene (20 ml) and evaporated to dryness. The residue was dissolved in dioxane (90 ml) and warmed to 60 °C, whereupon NaH (135 mg, 60% dispersion in mineral oil) was added. The solution was stirred for 15 minutes, then treated with 2-chloropropionic acid (675 µl) in dioxane (4.5 ml). The suspension was stirred for 2 hours at 70 °C, then treated portionwise with NaH (1.8 g) and dioxane (6 ml). The thick mixture was stirred at 50 °C for 16 hours, then treated cautiously with water (20 ml) to decompose excess NaH. The mixture was concentrated in vacuo to give a slurry which was diluted with water (15 ml) and washed with CHCl_3_ (15 ml). Ice and CHCl_3_ (45 ml each) were added, and the mixture treated with HCl (3M) until pH = 3. The layers were separated and the chloroform layer was further washed with ice-water (3 x 45 ml), dried, filtered, and concentrated *in vacuo*.

The crude product (140 mg) was dissolved in DMF (5 ml) and was treated with iodomethane (0.2 ml, 3.2 mmol) and K_2_CO_3_ (300 mg, 2.2 mmol). The mixture was stirred for 16 hours at room temperature, whereupon water and ethyl acetate (100 ml each) were added. The aqueous layer was further extracted with ethyl acetate (2 x 50 ml), and the combined organic layers were dried (Na_2_SO_4_), filtered, and concentrated *in vacuo*. The residue was purified by flash chromatography on silica gel (5% to 30% ethyl acetate in dichloromethane) to yield a smaller amount of the isomuramic acid derivative (see below) and the *title compound* (colourless solid, 44 mg, 85 µmol, 21%). ^1^H NMR (500 MHz, DMSO-*d*_6_) δ 8.20 (d, *J* = 9.2 Hz, 2H, 2 x ArH), 7.97 (d, *J* = 8.1 Hz, 1H, NH), 7.47 – 7.36 (m, 5H, 5 x ArH), 7.24 (d, J = 9.2 Hz, 2H, 2 x ArH), 5.72 (s, 1H, benzylidene CH), 5.47 (d, *J* = 7.7 Hz, 1H, H-1), 4.34 (q, *J* = 6.7 Hz, 1H, Lac-α), 4.29-4.25 (m, 1H, H-6a), 3.91 – 3.82 (m, 2H, H-2, H-3), 3.81 – 3.73 (m, 3H, H-4, H-5, H-6b), 3.61 (s, 3H, OMe), 1.81 (s, 3H, NAc), 1.27 (d, *J* = 6.7 Hz, 3H, Lac-Me). ^13^C NMR (126 MHz, DMSO-*d*_6_) δ 172.64 (C=O, <u>C</u>O_2_Me), 169.55 (N<u>C</u>OMe), 161.74 (C), 142.15 (C), 137.50 (C), 128.93 (CH), 128.28 (2 x CH), 125.90 (2 x CH), 125.86 (2 x CH), 116.67 (2 x CH), 100.15 (benzylidene CH), 98.05 (C-1), 80.82 (C-4), 77.71 (C-3), 75.43 (Lac-α), 67.60 (C-6), 65.64 (C-5), 54.29 (C-2), 51.76 (OMe), 23.01 (NCO<u>Me</u>), 18.75 (Lac-Me). HPLC-MS m/z: [M+H]^+^ Calcd for C_25_H_29_N_2_O_10_: 517.18; Found 517.15.

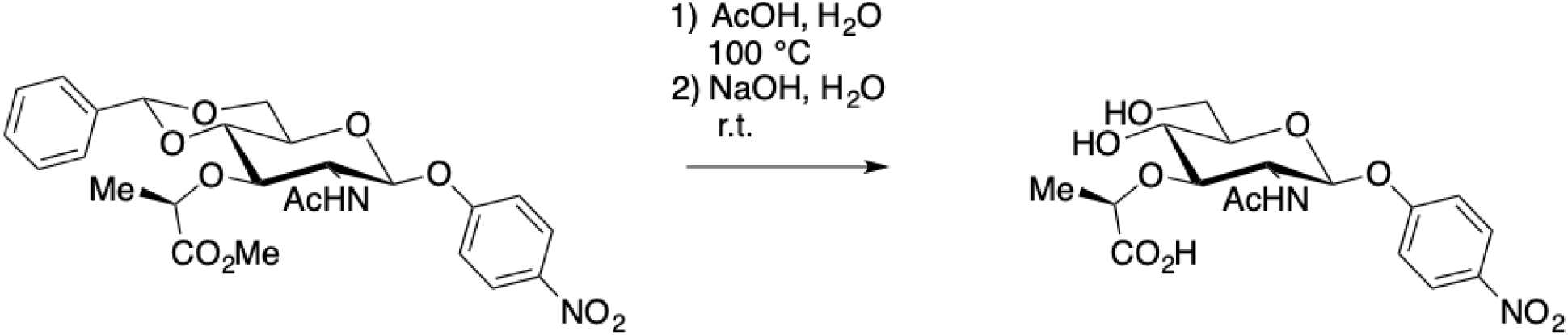

#### p-Nitrophenyl MurNAc

p-Nitrophenyl 3-O-(D-1-methoxycarbonylethyl)-4,6-benzylidine-GlcNAc (27 mg, 52 µmol) in 60% acetic acid (13 ml) was stirred at 100 °C for 45 minutes. The reaction mixture was concentrated *in vacuo*, diluted with sodium hydroxide solution (0.5 M, 5 ml), and stirred at room temperature for 1 hour. The reaction mixture was neutralized with Amberlite resin, filtered and lyophilized. The residue was purified by HPLC (5% to 95% acetonitrile in water + 0.1% formic acid) to yield the title compound (14.5 mg, 35 µmol, 67%) as a colorless solid. ^1^H NMR (300 MHz, D_2_O) δ 8.11 (d, *J* = 9.3 Hz, 2H, 2 x ArH), 7.06 (d, *J* = 9.3 Hz, 2H, 2 x ArH), 5.20 (d, *J* = 8.4 Hz, 1H, H-1), 4.32 (q, *J* = 6.8 Hz, 1H, Lac-α), 3.97 (t, *J* = 9.2 Hz, 1H, H-2), 3.91 – 3.83 (m, 1H, H-6a), 3.71 (dd, *J* = 12.3, 4.1, 1 H, H-6b) 3.66 – 3.48 (m, 3H, H-3, H-4, H-5), 1.89 (s, 3H, NHAc), 1.35 (d, *J* = 6.9 Hz, 3H, Lac-Me). ^13^C NMR (75 MHz, D_2_O) δ 177.83 (C=O), 175.26 (C=O), 162.32 (C), 143.22 (C), 126.69 (2 x CH), 117.10 (2 x CH), 99.16 (C-1), 83.14 (C-3), 78.20 (Lac-α), 76.67 (C-5), 69.87 (C-4), 61.02 (C-6), 55.22 (C-2), 22.84 (NHAc), 19.25 (Lac-Me). HRMS (ESI-TOF) m/z: [M+H]^+^ Calcd for C_17_H_23_N_2_O_10_: 415.1347; Found 415.1345.

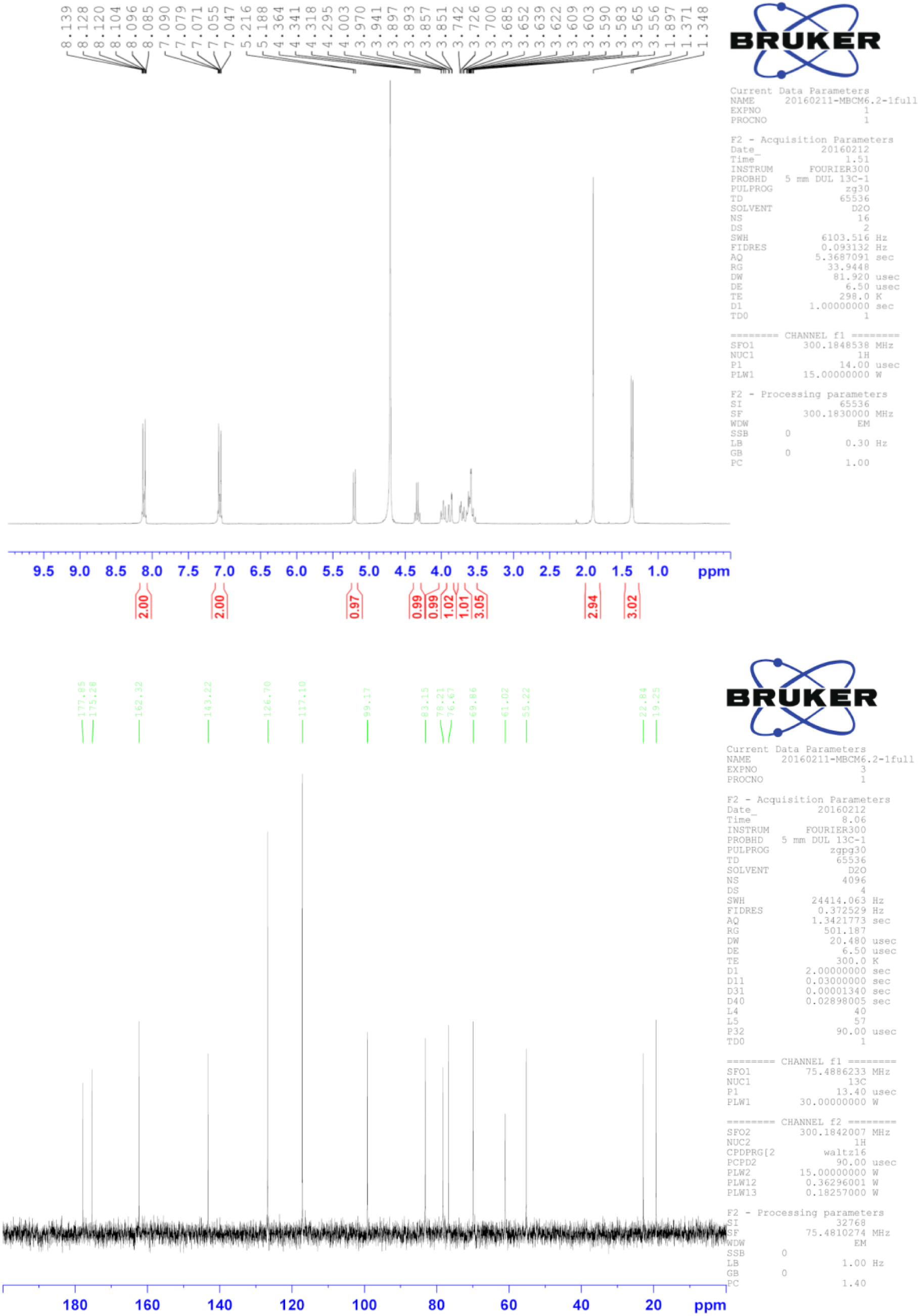

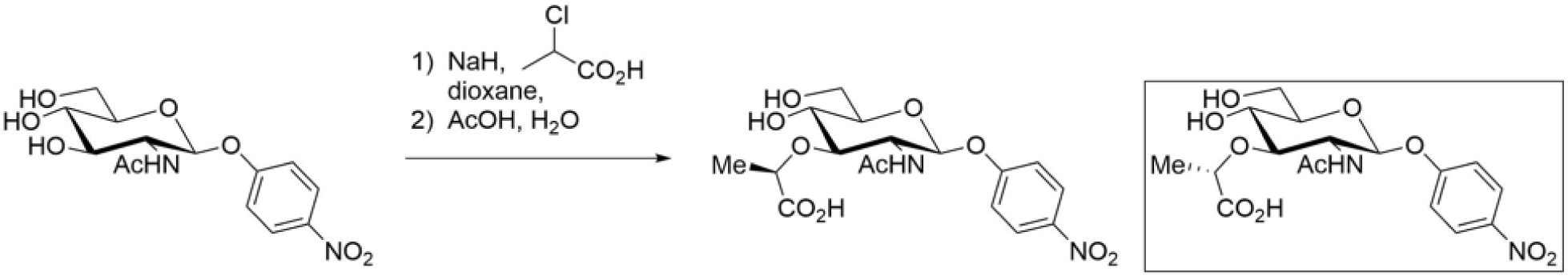

#### p-Nitrophenyl-isoMurNAc

p-Nitrophenyl 4,6-benzylidene-GlcNAc (57 mg, 0.13 mmol) was taken up in toluene (20 ml) and evaporated to dryness. The residue was dissolved in dioxane (30 ml) and warmed to 60 °C, whereupon NaH (45 mg, 60% dispersion in mineral oil) was added. The solution was stirred for 15 minutes, then treated with 2-chloropropionic acid (225 µl) in dioxane (1.5 ml). The suspension was stirred for 2 hours at 70 °C, then treated portionwise with NaH (0.6 g) and dioxane (2 ml). The thick mixture was stirred at 50 °C for 16 hours, then treated cautiously with water (20 ml) to decompose excess NaH. The mixture was concentrated in vacuo to give a slurry which was diluted with water (5 ml) and washed with CHCl3 (5 ml). Ice and CHCl3 (15 ml each) were added, and the mixture treated with HCl (3M) until pH = 3. The layers were separated and the chloroform layer was further washed with ice-water (3 x 15 ml), dried, filtered, and concentrated in vacuo to yield the crude product (50 mg).

A portion of the crude product (23 mg) was dissolved in 60% acetic acid (10 ml) and was stirred at 100 °C for 15 minutes. The reaction mixture was concentrated in vacuo and azeotroped with toluene. The residue was purified by HPLC (5% to 15% acetonitrile in water + 0.1% formic acid) to yield p-Nitrophenyl-MurNAc (2.9 mg, analytical data as described previously) and the title compound (1.7 mg, 4.1 µmol, 7%) as a colorless solid. 1H NMR (500 MHz, CD3OD) δ 8.26 (d, J = 9.3 Hz, 2H, 2 x ArH), 7.20 (d, J = 9.3 Hz, 2H, 2 x ArH), 5.34 (d, J = 8.5 Hz, 1H, H-1), 4.23 (q, J = 7.0 Hz, 1H, Lac-α), 4.12 (dd, J = 9.9, 8.5 Hz, 1H, H-2), 3.94 (dd, J = 12.6, 1.7 Hz, 1H, H-6a), 3.80 – 3.75 (m, 1H, H-6b), 3.70 – 3.60 (m, 3H, H-3, H-4, H-5), 2.05 (s, 3H, NHAc), 1.37 (d, J = 7.0 Hz, 3H, Lac-Me). 13C NMR (126 MHz, CD3OD) δ 180.29 (C=O), 175.40 (C=O), 162.25 (C), 143.36 (C), 126.74 (2 x CH), 117.19 (2 x CH), 98.81 (C-1), 83.27 (C-3), 78.90 (Lac-α), 76.79 (C-5), 69.90 (C-4), 60.99 (C-6), 55.37 (C-2), 22.67 (NHAc), 19.41 (Lac-Me). HRMS (ESI-TOF) m/z: [M+H]+ Calcd for C17H23N2O10: 415.1347; Found 415.1343

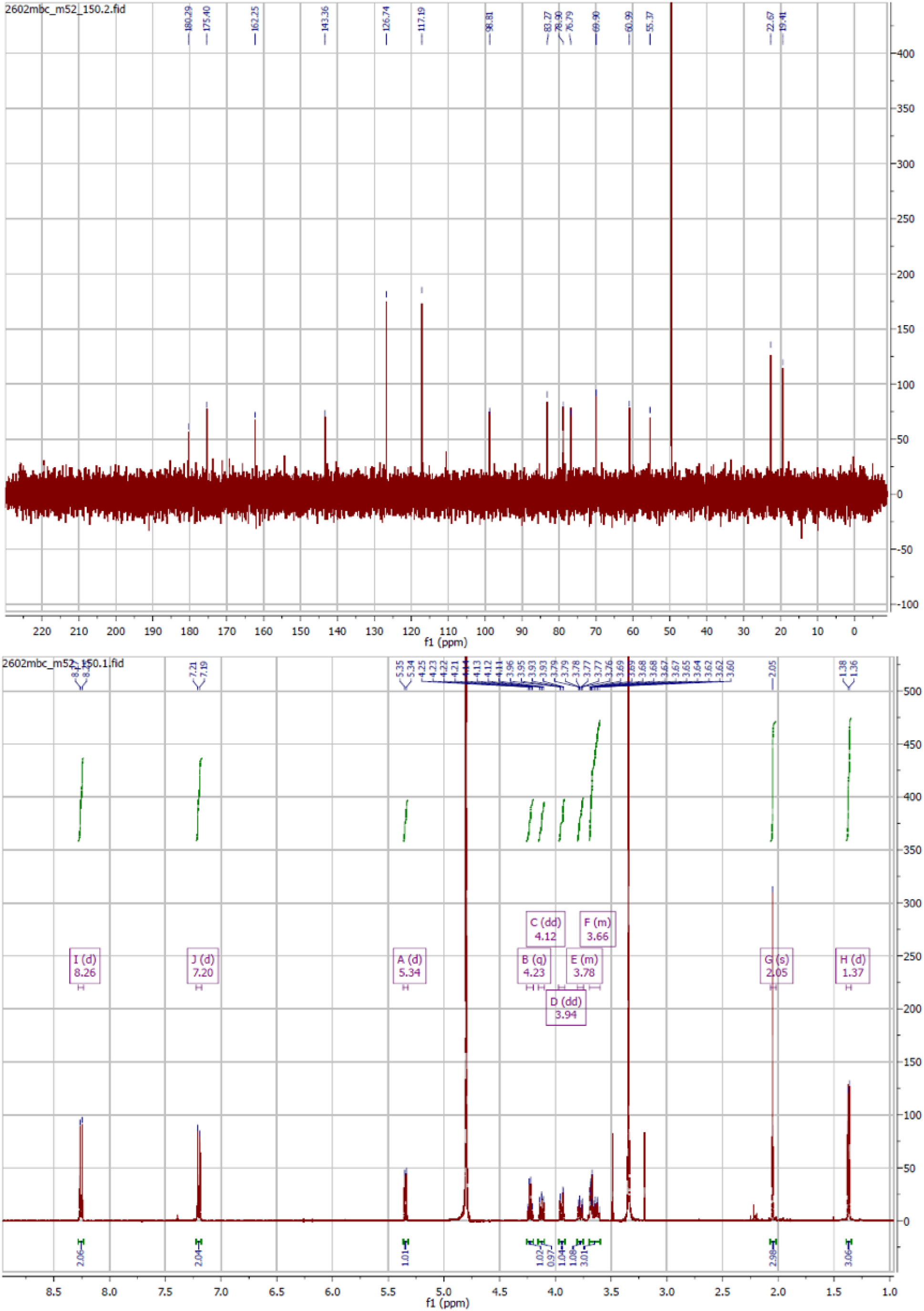

## Supporting Tables

**Table S1.**
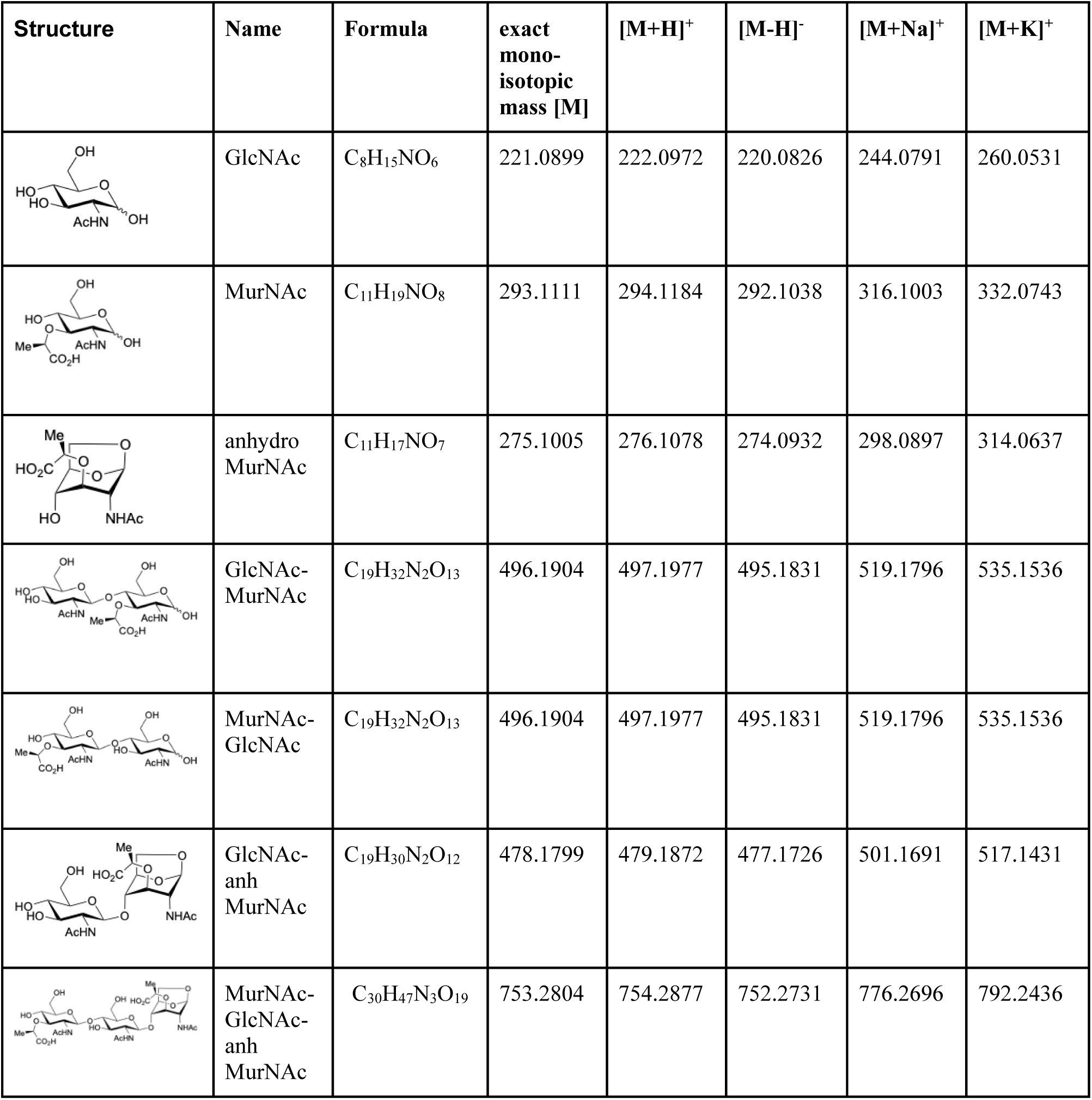
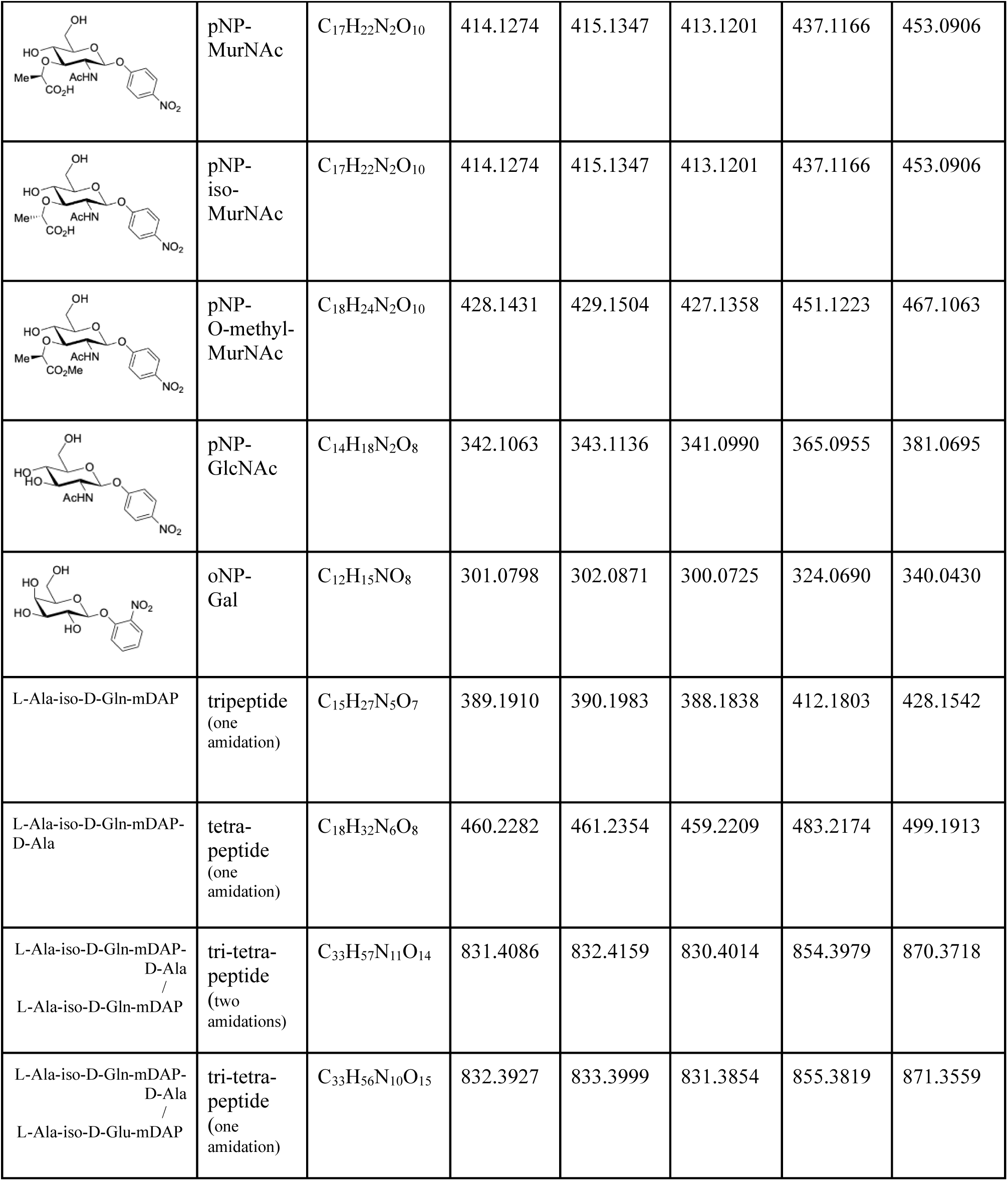
Structures and exact monoisotopic masses. Overview of peptidoglycan degradation products and chromogenic glycosidase substrates and their exact masses of proton, sodium, potassium adducts in positive ionization mode and the deprotonated species in negative ionization mode.

| Structure | Name | Formula | exact mono-isotopic mass [M] | [M+H]<sup>+</sup> | [M-H]<sup>-</sup> | [M+Na]<sup>+</sup> | [M+K]<sup>+</sup> |
| --- | --- | --- | --- | --- | --- | --- | --- |
|  | GlcNAc | C<sub>8</sub>H<sub>15</sub>NO<sub>6</sub> | 221.0899 | 222.0972 | 220.0826 | 244.0791 | 260.0531 |
|  | MurNAc | C<sub>11</sub>H<sub>19</sub>NO<sub>8</sub> | 293.1111 | 294.1184 | 292.1038 | 316.1003 | 332.0743 |
|  | anhydro MurNAc | C<sub>11</sub>H<sub>17</sub>NO<sub>7</sub> | 275.1005 | 276.1078 | 274.0932 | 298.0897 | 314.0637 |
|  | GlcNAc-MurNAc | C<sub>19</sub>H<sub>32</sub>N<sub>2</sub>O<sub>13</sub> | 496.1904 | 497.1977 | 495.1831 | 519.1796 | 535.1536 |
|  | MurNAc-GlcNAc | C<sub>19</sub>H<sub>32</sub>N<sub>2</sub>O<sub>13</sub> | 496.1904 | 497.1977 | 495.1831 | 519.1796 | 535.1536 |
|  | GlcNAc-anh MurNAc | C<sub>19</sub>H<sub>30</sub>N<sub>2</sub>O<sub>12</sub> | 478.1799 | 479.1872 | 477.1726 | 501.1691 | 517.1431 |
|  | MurNAc-GlcNAc-anh MurNAc | C<sub>30</sub>H<sub>47</sub>N<sub>3</sub>O<sub>19</sub> | 753.2804 | 754.2877 | 752.2731 | 776.2696 | 792.2436 |
| | pNP-MurNAc | $C_{17}H_{22}N_2O_{10}$ | 414.1274 | 415.1347 | 413.1201 | 437.1166 | 453.0906 |
| | pNP-iso-MurNAc | $C_{17}H_{22}N_2O_{10}$ | 414.1274 | 415.1347 | 413.1201 | 437.1166 | 453.0906 |
| | pNP-O-methyl-MurNAc | $C_{18}H_{24}N_2O_{10}$ | 428.1431 | 429.1504 | 427.1358 | 451.1223 | 467.1063 |
| | pNP-GlcNAc | $C_{14}H_{18}N_2O_8$ | 342.1063 | 343.1136 | 341.0990 | 365.0955 | 381.0695 |
| | oNP-Gal | $C_{12}H_{15}NO_8$ | 301.0798 | 302.0871 | 300.0725 | 324.0690 | 340.0430 |
| L-Ala-iso-D-Gln-mDAP | tripeptide<br>(one amidation) | $C_{15}H_{27}N_5O_7$ | 389.1910 | 390.1983 | 388.1838 | 412.1803 | 428.1542 |
| L-Ala-iso-D-Gln-mDAP-D-Ala | tetra-peptide<br>(one amidation) | $C_{18}H_{32}N_6O_8$ | 460.2282 | 461.2354 | 459.2209 | 483.2174 | 499.1913 |
| L-Ala-iso-D-Gln-mDAP-D-Ala<br>/<br>L-Ala-iso-D-Gln-mDAP | tri-tetra-peptide<br>(two amidations) | $C_{33}H_{57}N_{11}O_{14}$ | 831.4086 | 832.4159 | 830.4014 | 854.3979 | 870.3718 |
| L-Ala-iso-D-Gln-mDAP-D-Ala<br>/<br>L-Ala-iso-D-Glu-mDAP | tri-tetra-peptide<br>(one amidation) | $C_{33}H_{56}N_{10}O_{15}$ | 832.3927 | 833.3999 | 831.3854 | 855.3819 | 871.3559 |

**Table S2.**
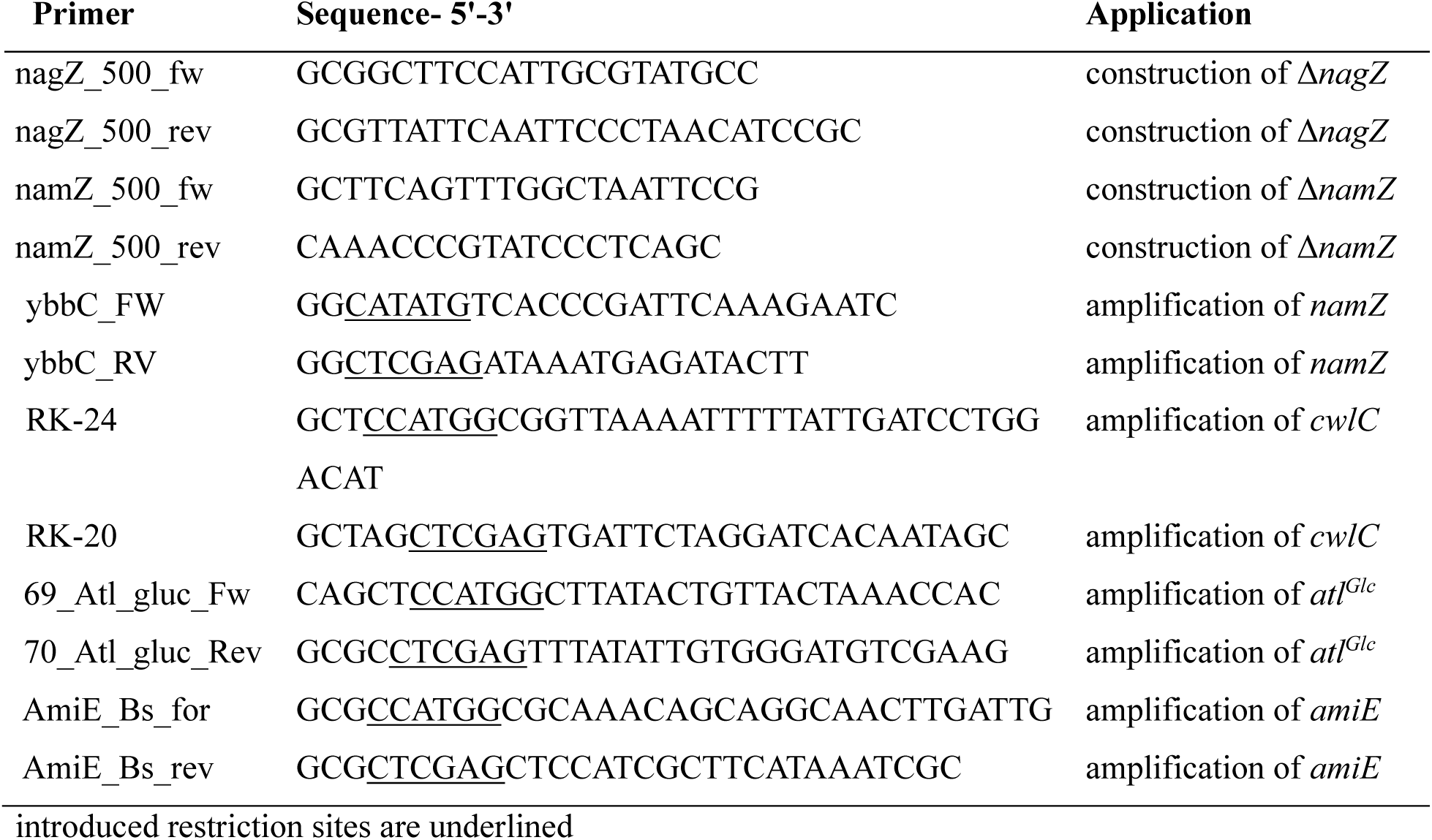
List of Oligonucleotides.

## Supporting Figures

**Figure S1.**
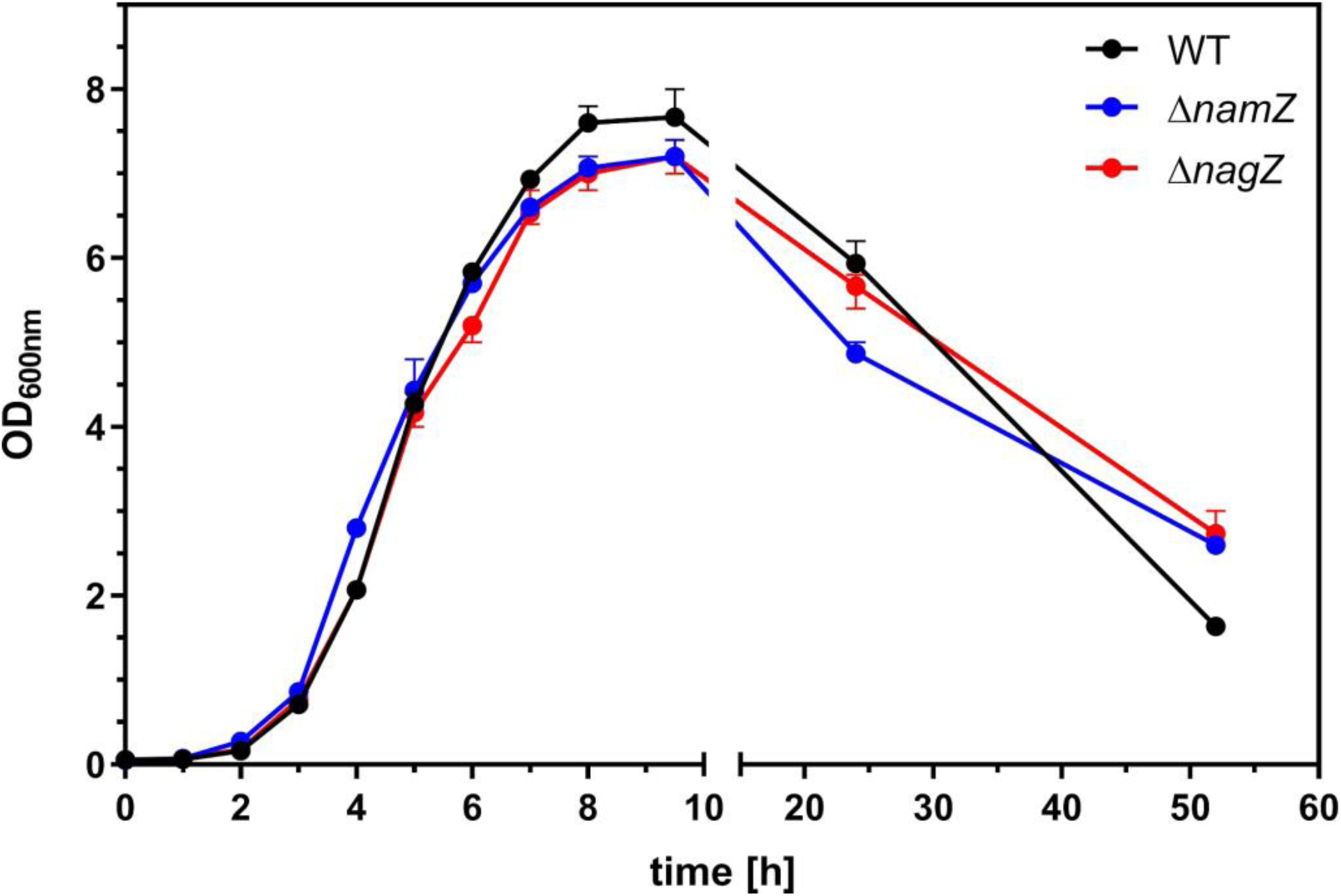
Comparison of the growth of *B. subtilis* WT, Δ*nagZ and* Δ*namZ* in LB medium under aerobic conditions. The Δ*nagZ* and Δ*namZ* mutants showed diminished growth in the transition to stationary phase (24 h) and an alleviated lytic behavior following prolonged stationary phase (52 h) when compared to WT. Cells were grown in 500 ml flasks with chicane in a final volume of 40 ml LB under constant shaking.

**Figure S2.**
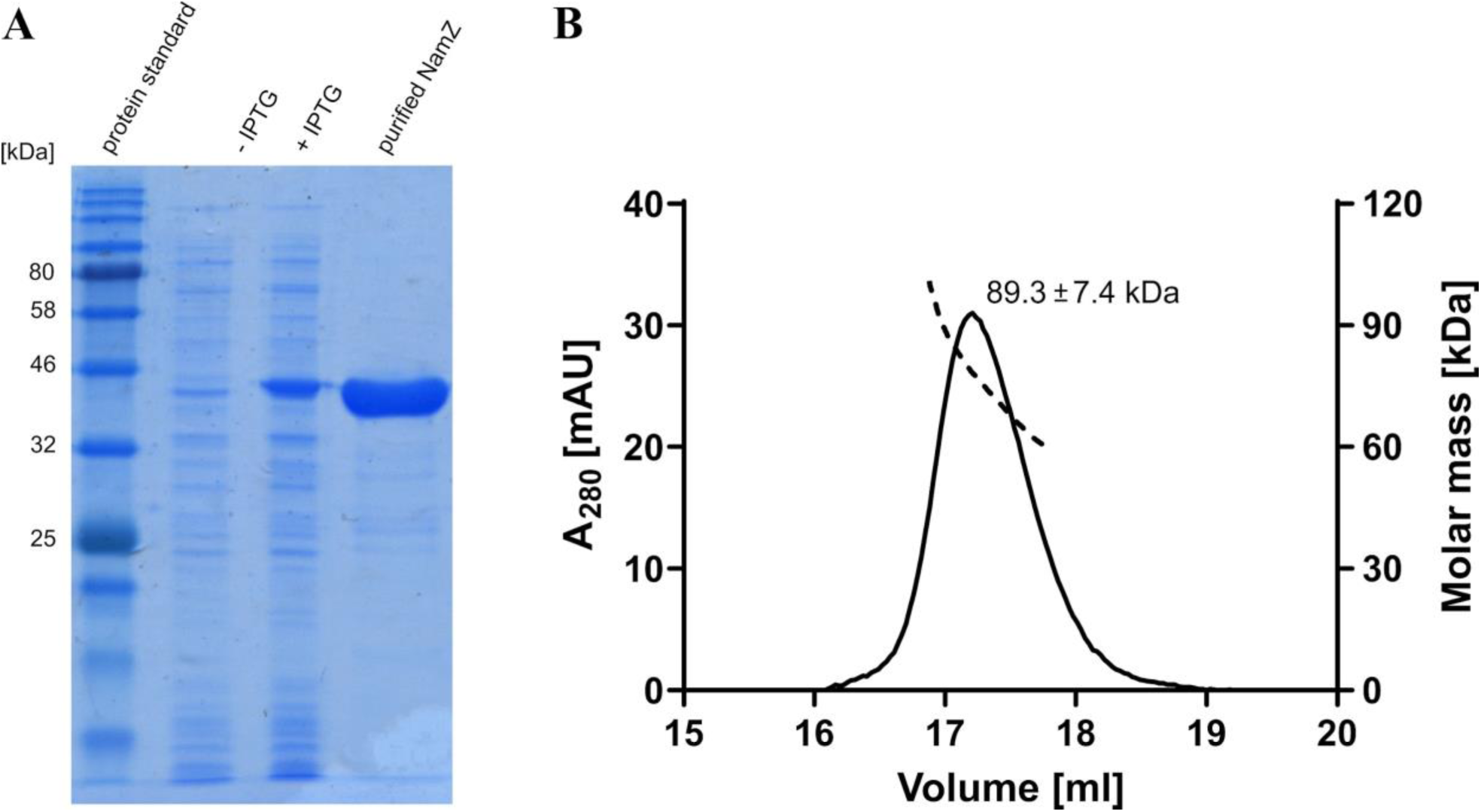
Purity of recombinant *B. subtilis* NamZ revealed by SDS-PAGE and identification of its dimeric state in solution by SEC-MALS analysis. **(A)** SDS-PAGE analysis of NamZ; lane 1, protein standard; lane 2, *E. coli* cell extract before induction (-IPTG) and, lane 3 after induction (+ IPTG); lane 4, pure recombinant NamZ protein after Ni^2+^ affinity and size exclusion chromatography (24 µg). The protein yield was 8 mg per L culture and appeared at a molecular mass consistent with the theoretical molecular mass of the monomer of 44.77 kDa. **(B)** SEC-MALS profile of NamZ. Recombinant NamZ was subject to a size-exclusion chromatography coupled with multiangle light scattering (SEC-MALS). The scattering data (dotted line) for the NamZ peak area was determined in three independent experiments resulting in an average of 89.3 ± 7.4 kDa at the peak maximum, which corresponds to the dimeric-mass of the monomer (44.77 kDa). The distribution of molar mass across NamZ peak was broad and ranged from ≈ 114.5 to 65.9 kDa, under native conditions.

**Figure S3.**
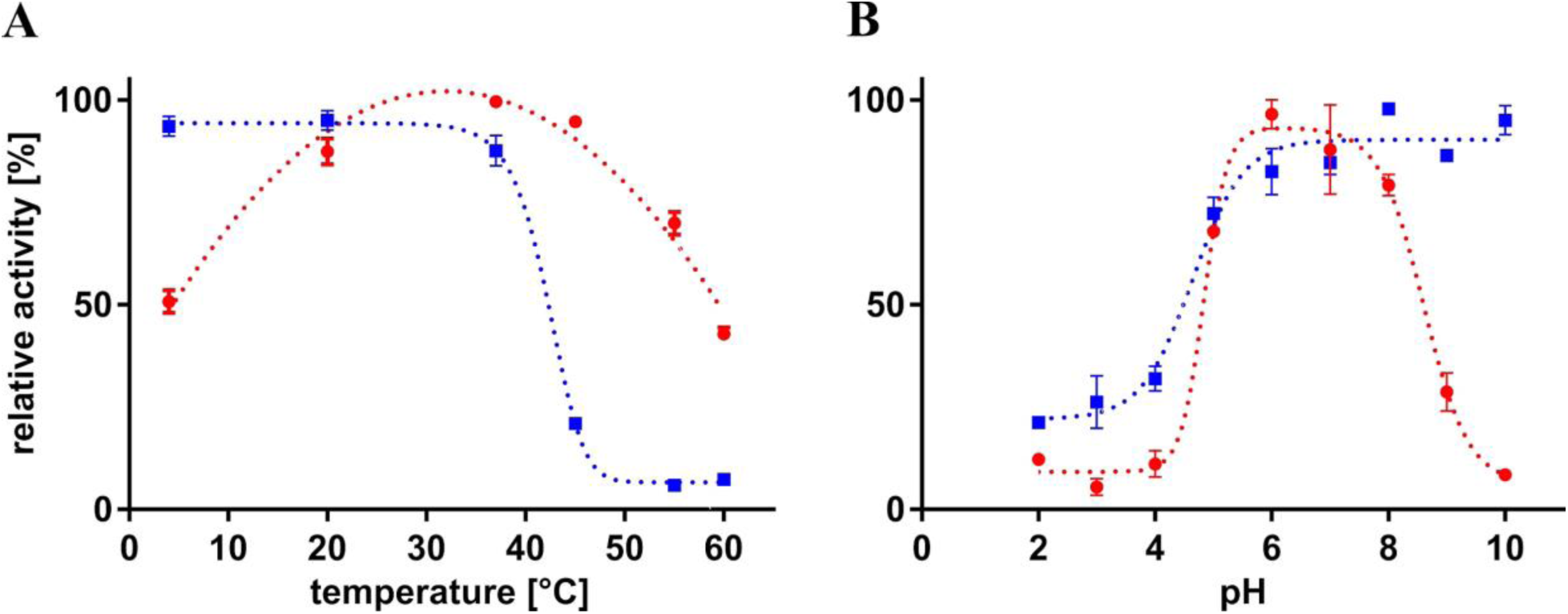
Temperature and pH stability and optima of NamZ. Enzyme stability (blue squares) and enzyme optimum activity (red circles) were assayed using pNP-MurNAc (0.2 mM) for 30 min at different **(A)** temperature and **(B)** pH conditions, using purified, recombinant NamZ (0.05 g/l). Standard errors (SEM) are indicated and calculated out of three biological replicates.

**Figure S4.**
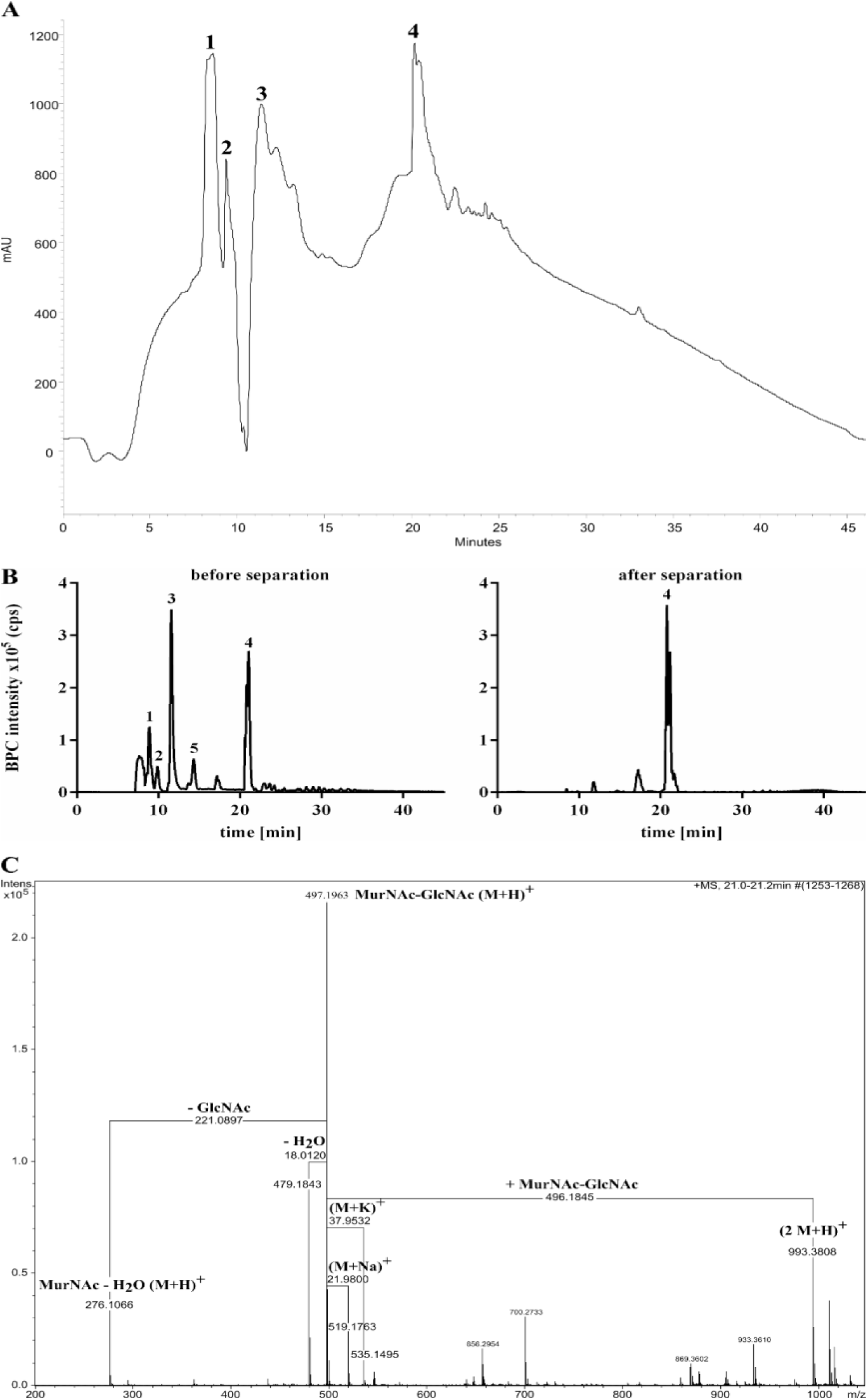
Separation from peptides and purification of MurNAc-GlcNAc by semi-preparative HPLC on a RP-C18 column. Peptidoglycan was digested overnight with CwlC and Atl^Glc^ and the reaction mixture was separated via HPLC **(A)**. The elution profile is shown at absorption of 202 nm. The main peaks (labeled 1-4) were identified by their m/z using LCMS **(B)**. Peaks 1-3 and 5 were identified as peptidoglycan-derived peptides: peak 1, at a retention time of 8.9 min, showed amidated tripeptide, peak 2, at a retention time of 9.9 min, showed amidated tetrapeptide, peak 3, at a retention time of 11.6 min, showed tri-tetrapeptide with two amidations, and peak 5, at a retention time of 14.3 min, showed tri-tetrapeptide with one amidation (Supporting Table S1). Peak 4, at a retention time of 21 min, solely contained MurNAc-GlcNAc. LCMS was also used to control the purity of the MurNAc-GlcNAc pool before (left) and after (right) purification by HPLC. Results are presented as base peak chromatograms (BPC). The mass spectrum of the MurNAc-GlcNAc pool after separation (peak 4; B) was further analyzed for purity by LCMS **(C)**. Main mass was (M+H)^+^ 497.196 m/z consistent with the theoretical mass of MurNAc-GlcNAc of (M+H)^+^ 497.1977 m/z). All other masses also could be assigned to MurNAc-GlcNAc: a dimer (2M+H)^+^ of MurNAc-GlcNAc, sodium and potassium ion adducts, as well as water and MurNAc elimination products.

**Figure S5.**
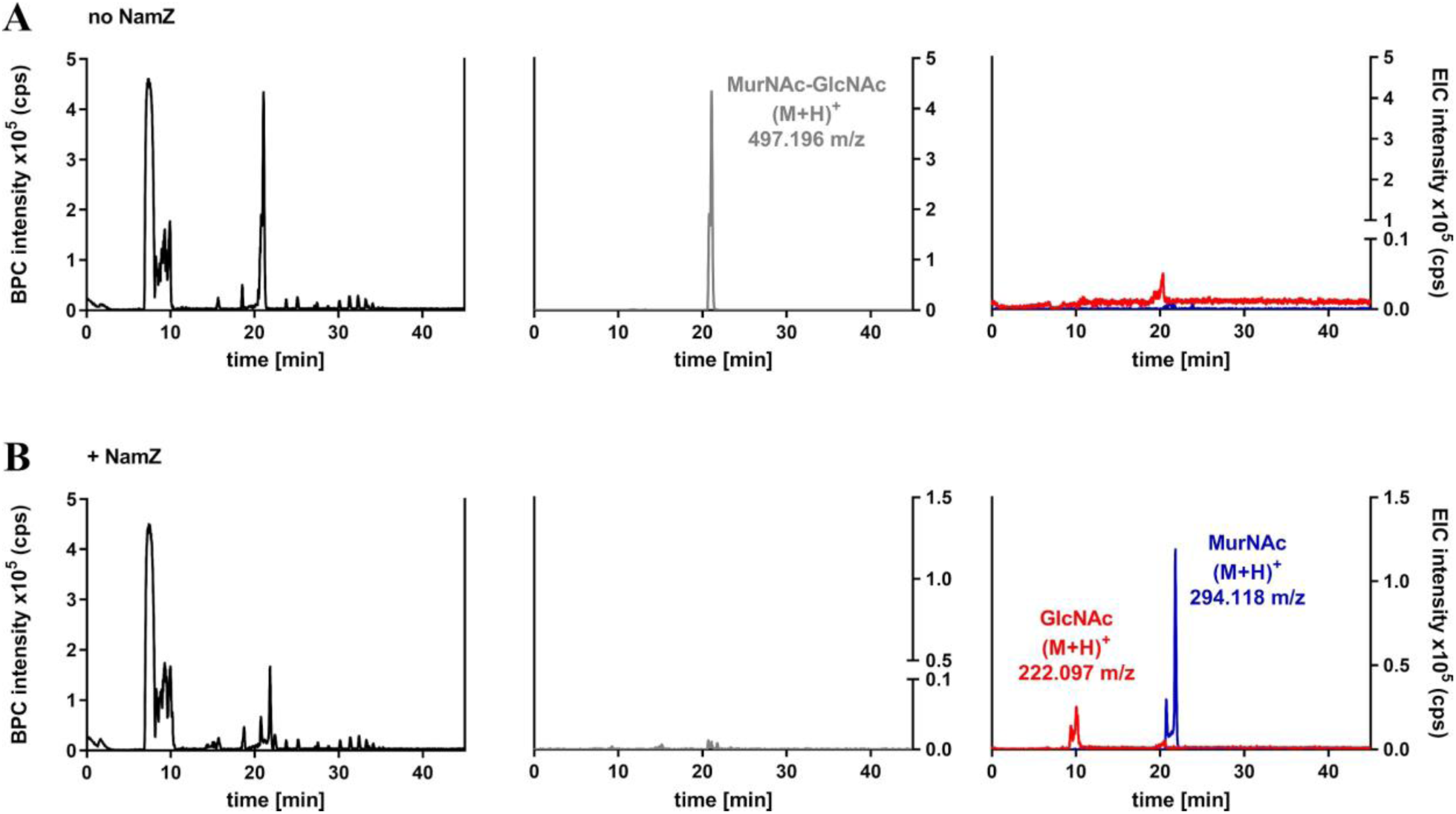
NamZ hydrolyzes MurNAc-GlcNAc yielding MurNAc and GlcNAc. Purified MurNAc-GlcNAc (0.1 mM) was incubated for 30 minutes at 37°C with NamZ (20 ng). Samples were analyzed by LC-MS in positive ion mode. Data are shown as base peak chromatograms (BPC; black, left y-axis) and extracted ion chromatograms (EIC; right y-axis). For MurNAc-GlcNAc without NamZ **(A)** a peak corresponding to MurNAc-GlcNAc at a retention time of 21.1 min with an observed mass of (M+H)^+^ 497.196 m/z could be detected (grey). For MurNAc-GlcNAc incubated with NamZ **(B)** two peaks could be detected: one for GlcNAc at a retention time of 10.0 min and with an observed mass of (M+H)^+^ 222.097 m/z (red) and the other corresponding to MurNAc at a retention time of 21.8 min and an observed mass of (M+H)^+^ 294.118 m/z (blue).

**Figure S6.**
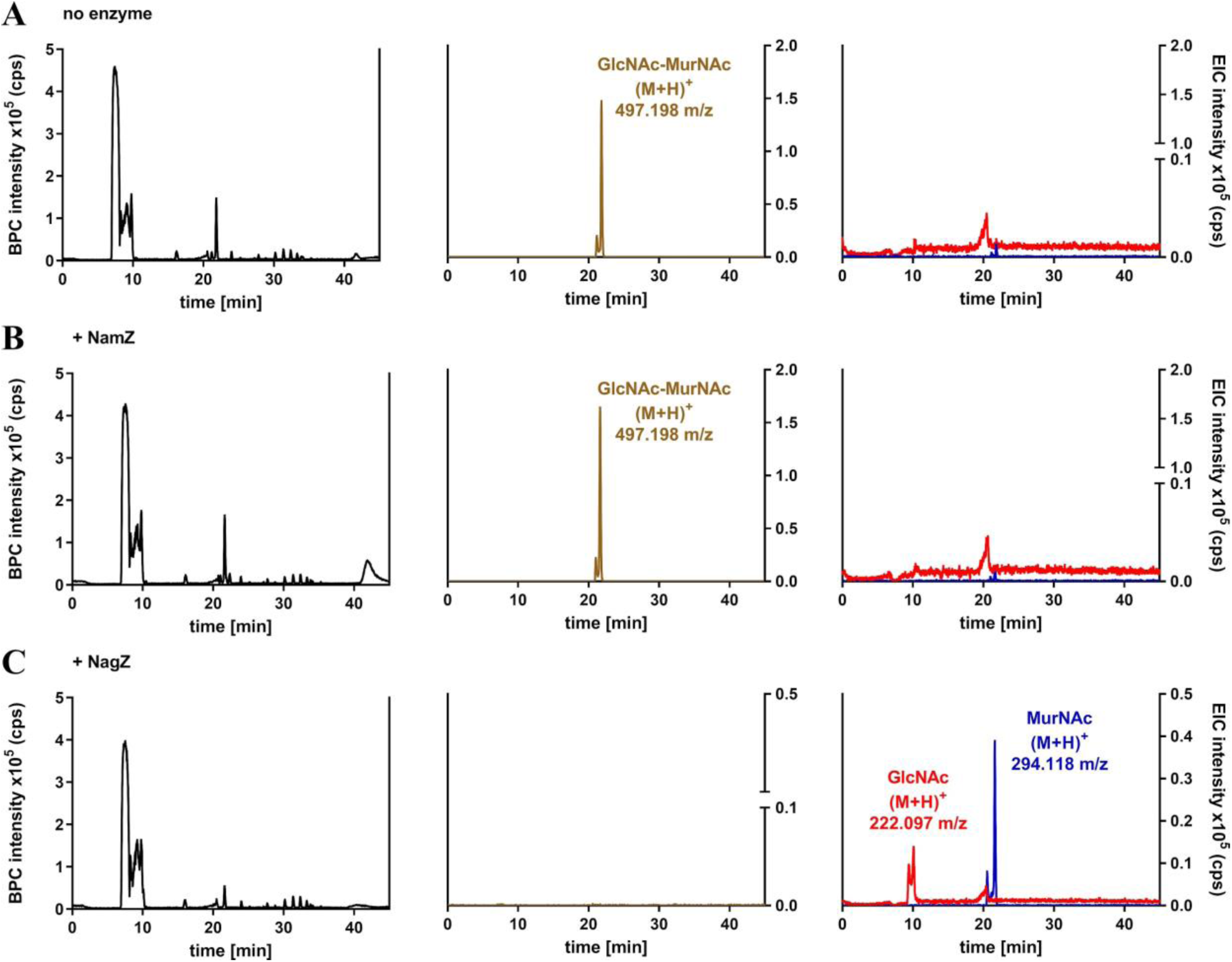
NamZ specifically requires MurNAc at the non-reducing end of a disaccharide. GlcNAc-MurNAc (0.1 mM) was incubated for 30 minutes at 37°C with either NamZ (+ NamZ) or NagZ (+NagZ) (each 20 ng), respectively. Samples were analyzed by LC-MS. Data are shown in base peak chromatograms (BPC; black, left y-axis) extracted ion chromatograms (EIC; right y-axis). For GlcNAc-MurNAc without NamZ **(A)** a peak corresponding to GlcNAc-MurNAc at a retention time of 21.8 min with an observed mass of (M+H)^+^ 497.198 m/z could be detected (brown). For GlcNAc-MurNAc with NamZ **(B)** a peak corresponding to GlcNAc-MurNAc at a retention time of 21.6 min with an observed mass of (M+H)^+^ 497.198 m/z could be detected (brown). For GlcNAc-MurNAc with NagZ **(C)** a peak corresponding to GlcNAc at a retention time of 10.1 min with an observed mass of (M+H)^+^ 222.097 m/z (red) and a peak corresponding to MurNAc at a retention time of 21.6 min with a mass of (M+H)^+^ 294.118 m/z (blue) could be detected.

**Figure S7.**
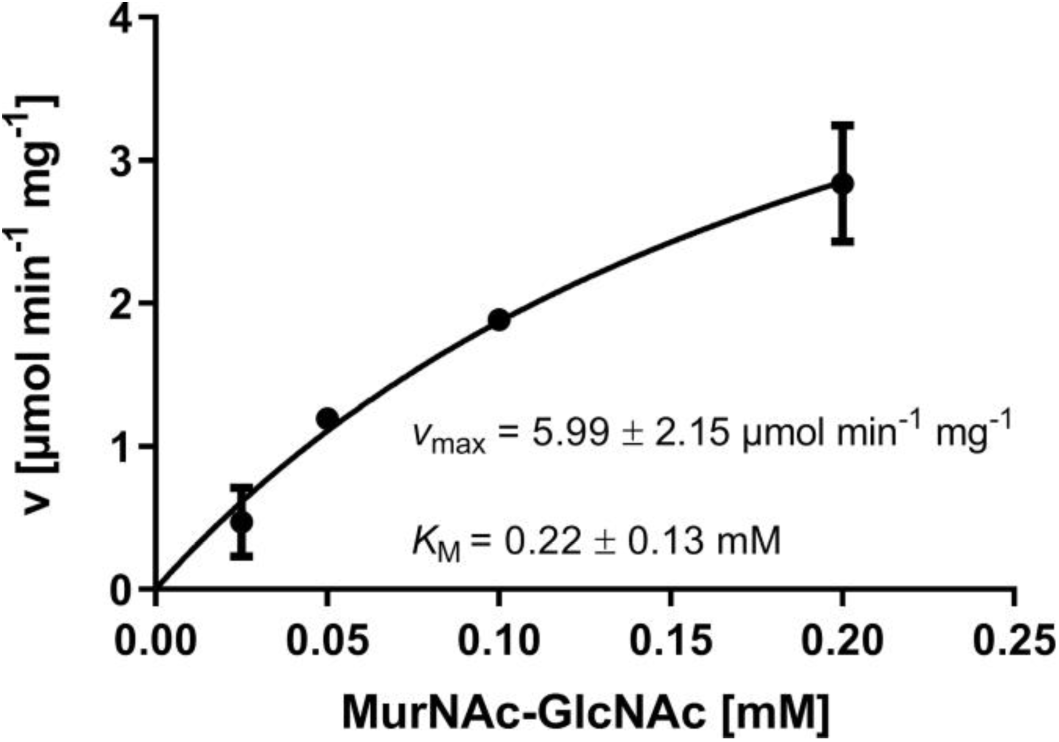
Kinetic parameters of NamZ determined for lower concentrations of MurNAc-GlcNAc substrate at pH 7.0. Kinetic parameters using purified MurNAc-GlcNAc as substrate were determined for substrate concentrations from 0.025 to 0.2 mM as described in Experimental Procedures. Standard errors (SEM) are indicated and calculated out of three biological replicates.

**Figure S9.**
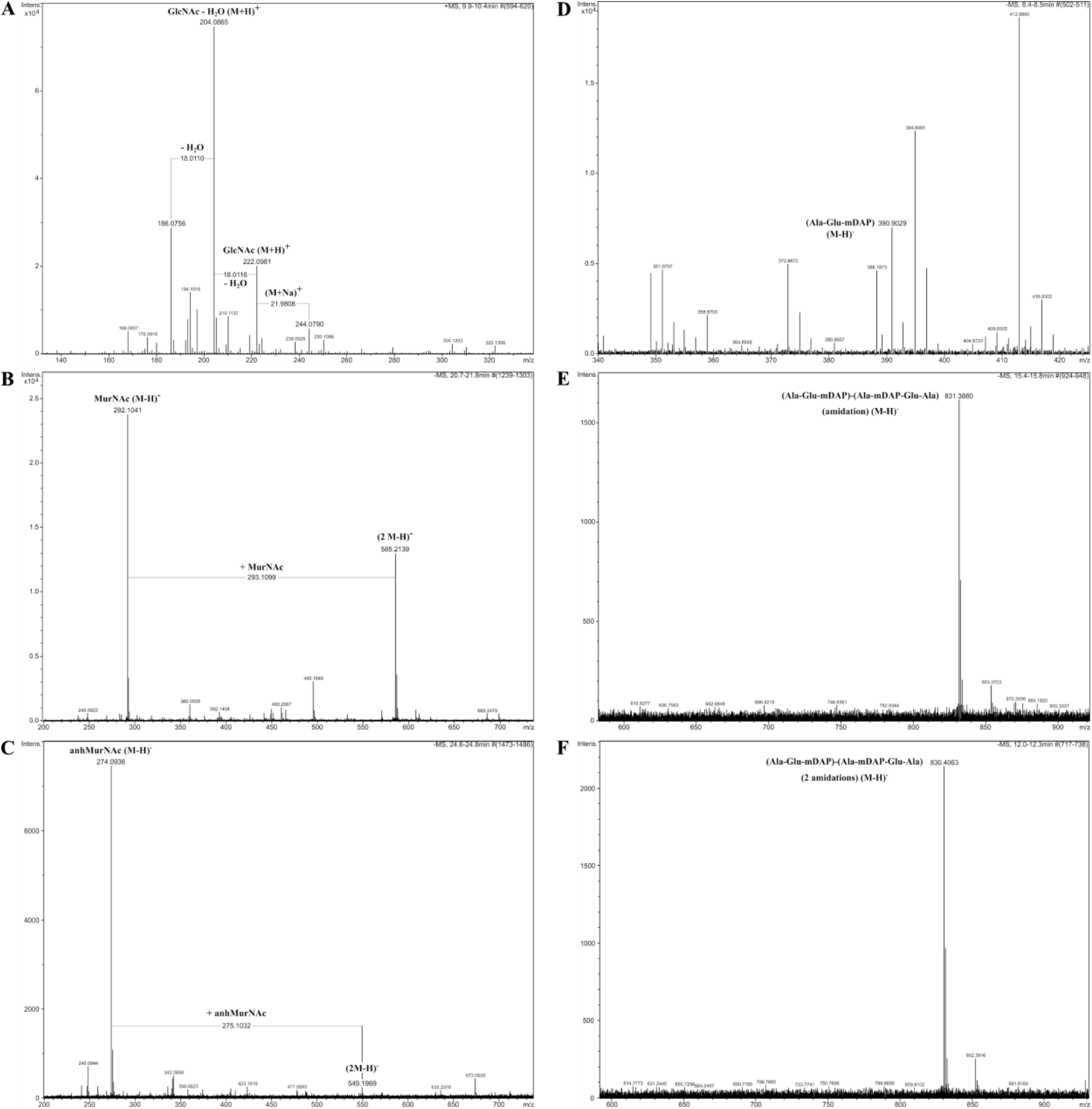
Sequential digest of peptidoglycan with NagZ, AmiE and NamZ of *B. subtilis*. Purified peptidoglycan was sequentially digested by the action of the three recombinant enzymes and analyzed by HPLC-MS. Measurement was performed in both positive and negative ion mode. Shown are the mass spectra of released peptidoglycan carbohydrates (A – C) and peptides (D – F). **(A)** GlcNAc release was identified by the appearance of the mass of (M+H)^+^ 222.098 m/z. Also masses for the sodium adduct ((M+H)^+^ 244.079 m/z) and water elimination products appeared. **(B)** MurNAc release was identified by the appearance of a mass of (M-H)^-^ 292.104 m/z. Also the mass for MurNAc dimer ((2 M-H)^-^ 585.214 m/z) appeared. **(C)** AnhMurNAc could be identified by the appearance of a mass of ((M-H)^-^ 274.094 m/z). Also the mass for anhMurNAc dimer ((2 M-H)^-^ 549.197 m/z), which occurs due to ionization conditions. **(D)** Tripeptide with an amidation could be identified with a mass of (M-H)^-^ 388.187 m/z. **(E)** Tri-tetrapeptide with an amidation could be identified with a mass of (M-H)^-^ 831.388 m/z. **(F)** Tri-tetrapeptide with two amidations could be identified with a mass of (M-H)^-^ 830.406. See Supporting Table S1 for theoretical m/z values.

